# snCED-seq: High-fidelity cryogenic enzymatic dissociation of nuclei for single-nucleus RNA-seq of FFPE tissues

**DOI:** 10.1101/2024.09.20.614201

**Authors:** Yunxia Guo, Junjie Ma, Ruicheng Qi, Xiaoying Ma, Jitao Xu, Kaiqiang Ye, Yan Huang, Xi Yang, Guang-zhong Wang, Xiangwei Zhao

## Abstract

Profiling cellular heterogeneity in formalin-fixed paraffin-embedded (FFPE) tissues is key to characterizing clinical specimens for biomarkers, therapeutic targets, and drug responses. Recent advancements in single-nucleus RNA sequencing (snRNA-seq) techniques tailored for FFPE tissues have demonstrated their feasibility. However, isolation of high-quality nuclei from FFPE tissue with current methods remains challenging due to RNA cross-linking. We, therefore, proposed a novel strategy for the preparation of high-fidelity nuclei from FFPE samples, cryogenic enzymatic dissociation (CED) method, and performed snRandom-seq (snCED-seq) for polyformaldehyde (PFA)-fixed and FFPE brains to verify its applicability. The method is compatible with both PFA-based and FFPE brains or other organs with less hands-on time and lower reagent costs, and produced 10 times more nuclei than the homogenate method, without secondary degradation of RNA, and maximized the retention of RNA molecules within nuclei. snCED-seq shows 1.5-2 times gene and UMI numbers per nucleus, higher gene detection sensitivity and RNA coverage, and a minor rate of mitochondrial and ribosomal genes, compared with the nuclei from traditional method. The correlation gene expression of nucleus from the post-fixed and the frozen sample can be up to 94 %, and the gene expression of our nuclei was more abundant. Moreover, we applied snCED-seq to cellular heterogeneity study of the specimen on Alzheimer’s Disease (AD) to demonstrate a pilot application. Scarce Cajal Retzius cells in older mice were robustly detected in our data, and we successfully identified two subpopulations of disease-associated in astrocytes, microglia and oligodendrocytes, respectively. Meanwhile, we found that most cell types are affected at the transcriptional level by AD pathology, and there is a disease susceptibility gene set that affects these cell types similarly. Our method provides powerful nuclei for snRNA-seq studies for FFPE specimens, and even helps to reveal multi-omics information of clinical samples.

## 1. Introduction

High-throughput single-cell/nuclei RNA sequencing (scRNA/snRNA-seq) methods have revolutionized the entire-field of biomedical research [1–3]. scRNA/snRNA-seq have been highly successful at disease mechanisms, discovering biomarkers to help stratify patients, and identifying novel therapeutic targets as well as determining the impact of drugs. However, fresh/frozen specimen procurement is not a standard clinical and diagnostic practise in most institutions, and fresh/frozen samples cannot be obtained for certain sample types. Routine formalin-fixed paraffin-embedded (FFPE) tissues are the most common archivable specimens, constituting a vast and valuable patient material bank for clinical history [4]. Inevitably, the irreversible modifications caused by formalin fixation on macromolecules in FFPE samples always make it challenging for molecular biology applications. The studies have made great progress in transcription profiling in FFPE samples by optimal RNA extraction methods [5, 6] or spatial in situ profiling [7]. What’s more, the combinations of scRNA-seq and spatial technologies have been applied to FFPE tissues [3]. Currently, three methods have been posted, snPATHO-Seq [8], snFFPE-seq [9], and snRandom-seq [10], provided optimized methods to isolate single intact nuclei from FFPE tissues to perform snRNA-seq, which demonstrates the feasibility of snRNA-Seq in FFPE tissues and unlocks a dimension of these hard-to-use samples. Accurate transcriptomic characterization of each cell in clinical FFPE specimens is believed to provide a better understanding of cell heterogeneity and population dynamics, thereby improving accurate diagnosis, treatment, and prognosis of human disease. With the development of snRNA-seq techniques for FFPE samples, there is growing interest in the use of the vast archives of samples for diagnostic purposes.

The application of snRNA-seq in FFPE samples is premised on obtaining superior nuclei. However, isolation of intact and high-quality nuclei remains challenging due to RNA crosslinking, modification, and degradation caused by formaldehyde fixation. The strategies of nuclei preparation for FFPE tissues are longstanding and can date back to the last century, but previous applications are only limited to DNA content [11], fluorescence in situ hybridization (FISH) [12, 13], genome-wide association studies (GWAS) [14, 15], and chromatin accessibility profiling [16]. Specifically, nuclei were dissociated by hyperthermia of biological tissue sections in protease solution, a technique that is sensitive to heating time and easily destroys the nuclear membrane, resulting in the loss of nuclear morphology. Moreover, prolonged exposure to enzyme buffers may increase the permeability of the nuclear membrane, resulting in RNA molecule leakage and adversely affecting snRNA-seq experiments conducted in droplets. The current state-of-the-art snRNA-seq for FFPE samples uses a mechanical homogenization method that is suitable for frozen samples before, and combined with a hyperthermic enzyme dissociation approach for nuclei preparation [8–10]. However, the homogenization of formaldehyde-fixed tissue poses challenges, leading to the presence of debris in the resulting nuclei suspension, which necessitates multiple filtration steps. This, in turn, affects the yield of nuclei and may result in the loss of smaller nuclei. However, the presence of tissue debris remains a challenge, introducing a higher amount of ribosomal RNA (rRNA), which can affect sequencing data quality. Therefore, the acquisition of high-quality nuclei from PFA-fixed or FFPE samples will be an important basis for transcriptome study of clinical samples.

To overcome these challenges, we proposed CED method, an efficient and high-fidelity method to extract nuclei from FFPE tissues, and combine it with full-length and total RNA snRNA sequencing for post-fixed brains. We reported side-by-side comparison of nuclei between CED and conventional methods, as well as among fresh frozen, PFA-fixed and FFPE tissues, to validate the robustness of snCED-seq in FFPE samples. Although we have optimized the conventional method, the nuclei obtained by CED method outperformed those reported method in terms of RNA integrity, nuclei numbers, number of gene and UMI per nuclei and richness of gene expression. Next, we used snCED-seq to more than 60,000 single nuclei from AD hippocampus, and our resource provides a comprehensive cellular heterogeneity analysis of AD mice by characterizing the total transcriptome at single-cell resolution. We provided evidence to show that snCED-seq is a reliable platform for analysis the transcriptomic profiles of FFPE tissues of degenerative neurological diseases, and we believe that it also has the potential to unlock the largely untapped other archives of biological material found in pathology archives, paving the way for clinical applications.

## 2. Results

### 2.1 Overview of the cryogenic enzymatic dissociation of nuclei for post-fixed tissues

The acquisition of high-fidelity nuclei is a prerequisite for the research and application of snRNA-seq for FFPE samples, and also a key factor for its full mining of transcriptional information. Since the last century, nuclear preparation of FFPE tissues requires enzymatic dissociation at high temperatures (HED) worldwide and are limited to non-transcriptomic applications. We converted the idea of traditional protocols of preparing nuclei from FFPE tissues, the factors that we deemed pertinent to affect transcriptome analysis, such as dissociation temperature, reagent and time. We established a completely new method of nucleus preparation for post-fixed (paraformaldehyde fixed (PFA-fixed) and FFPE) tissues, cryogenic enzymatic dissociation (CED) strategy. For this method, sarcosyl was used instead of SDS (Sodium n-Dodecyl Sulfate) or Triton X-100 as an anionic surfactant to participate in the nuclei preparation, which was more friendly to the nuclear membrane than the cell membrane, and became the preferred component for nucleus isolation in CED method. Moreover, proteinase K (PK) was used to digest proteins of tissue to minimize background contamination. Our CED method eliminates the need for ultracentrifugation through a sucrose cushion, nor any filtration procedure, maximizing product retention, increasing nucleation rates, and free of nuclear membranes and cytoplasmic contamination. Most importantly, the entire nucleus preparation process was carried out at low temperature, which protects the nuclear membrane and maximally retains the RNA molecules within nuclues, providing high-fidelity nucleus for snRNA-seq research of FFPE samples. In addition, by adjusting the experimental parameters, CED method is not only suitable for tissue slides, but also has good compatibility with FFPE blocks, which is more in line with the application needs of snRNA-seq in disease research. Next, the full-length total RNAs within nuclei of frozen, PFA-fixed and FFPE brains were captured by random primers for snRNA-seq (snCED-seq), and the main workflow of snCED-seq was shown in Fig. 1.

**Fig. 1.**
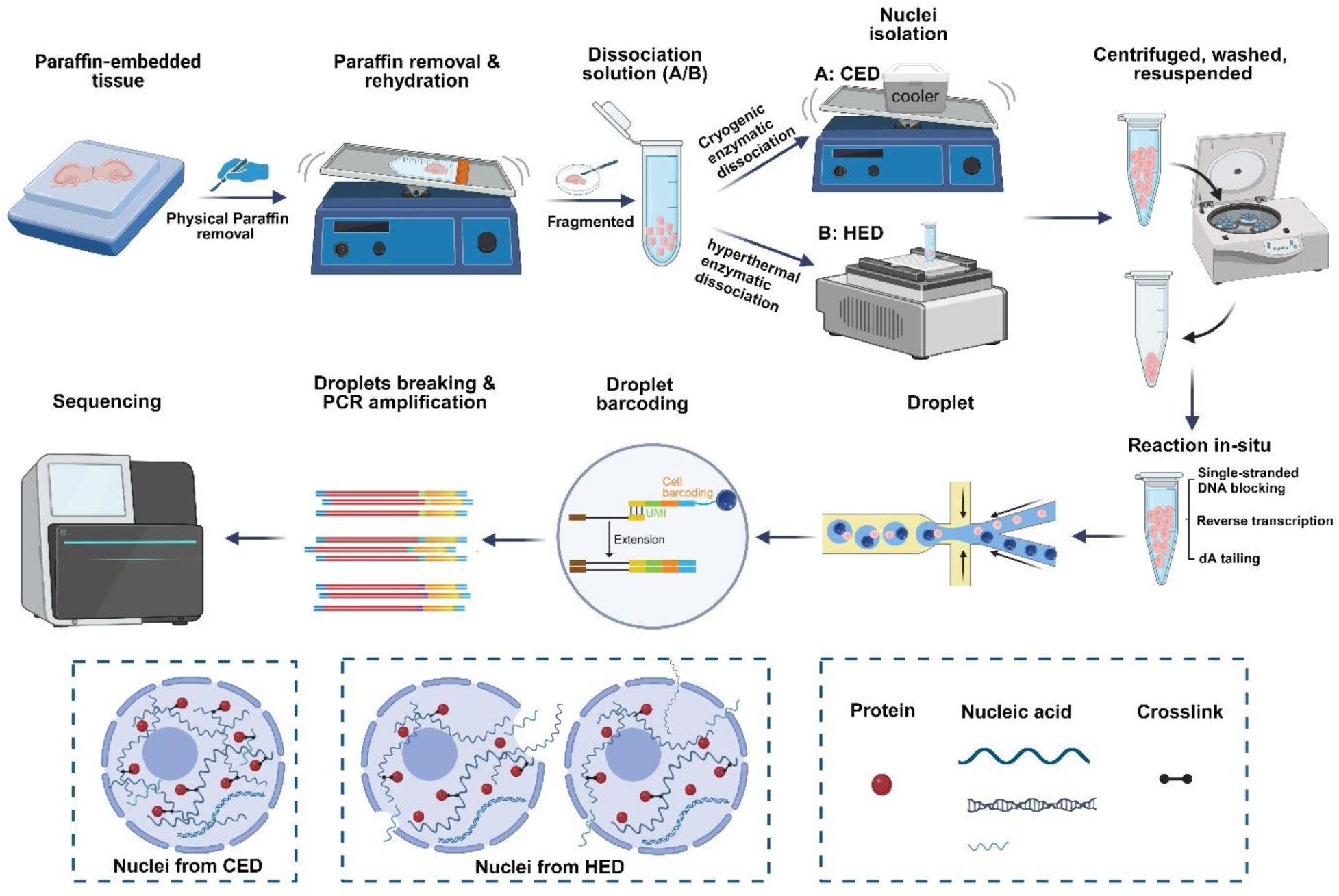
snCED-seq for post-fixed tissues overview. The workflow of snCED-seq for post-fixed tissues includes single nuclei isolation by CED and HED method with the snRandom-seq method used in this study. The steps from nucleus extraction to targeted sequencing are shown. In contrast to HED, the nuclei prepared with CED were morphologically intact without leakage of RNA molecular.

The nuclei drived from FFPE brains prepared by CED method had intact morphology, good dispersion, high purity without agglomeration (Fig. 2A). Confirmation of the integrity and dispersion of nuclear morphology was also verified using epifluorescence microscopy (Fig. 2B). Representative images of nuclei isolated from the hippocampus of three biological replicates showed much less debris and a size-distributions were centered around 6-8 µm (Fig. S1A), slightly smaller than normal frozen brain nuclei [17], presumably due to the tissue being fixed. Perhaps, CED method without cumbersome filtering procedures, tiny nuclei could be preserved. Statistics showed that at least a million levels of nuclei were obtained from each pair of hippocampi (Fig. S1A, bottom). The recent snRNA-seq techniques for FFPE tissues based on random primer capture [10] or gene probe capture [8] require the input of nearly one million nuclei to ensuring the output of about 10,000 nuclei. Our CED method strategy can effectively circumvent the shortcomings of the current two mainstream nuclear preparation strategies, and can export nuclei stably without introducing more impurities and destroying the nuclear membrane.

**Fig. 2.**
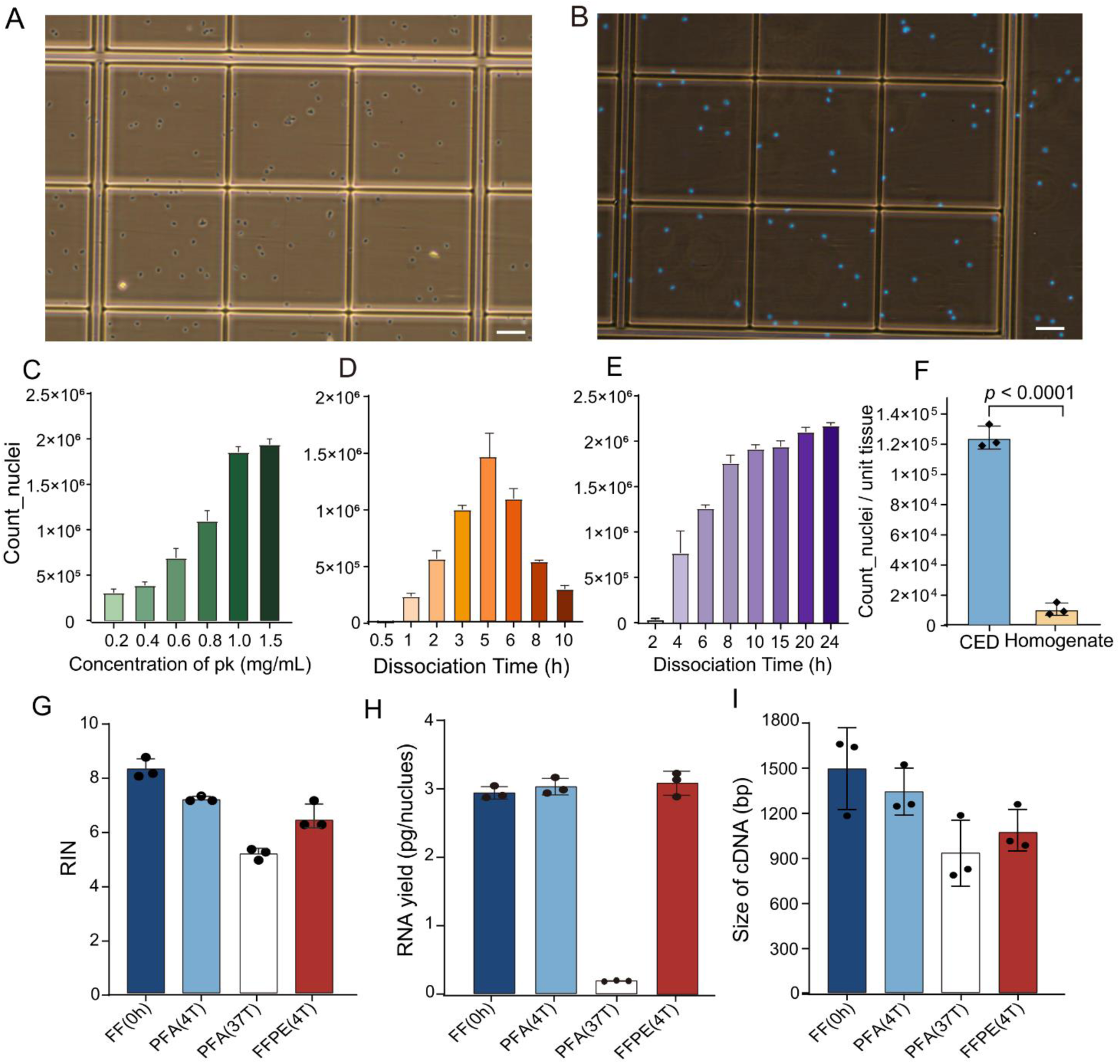
Quality control of nuclei prepared by cryogenic enzymatic dissociation. (**A, B**) Image of trypan blue-stained (A) and DAPI-stained (B) nuclei isolated from FFPE mouse brain by CED before cell encapsulation, respectively. Scale bar, 50 μm; (**C**) Nuclei yield at different proteinase K concentrations; (**D, E**) Bar plots showing the relationship between the nuclei numbers of the PFA-fixed hippocampus with the dissociation time was isolated by HED (C) and CED (D) methods; (**F**) Nuclear yield per gram of hippocampal tissue, using CED and mechanical homogenization, respectively; (**G, H**) RNA integrity number (RIN) (F) and total RNA yield of RNA extracted from nuclei of three forms of tissue; (**I**) Representative peak values of amplified cDNA in different groups. n = 3 technical replicates, and bars show mean ± s.d (C-H).

We optimized the proteinase K (PK) concentration in the nuclear dissociation system as measured by morphology and count of nuclei, and found that the optimal concentration for HED was 0.4 mg/mL, while higher was required in the CED for mouse brain, which was due to the reduced enzyme activity at low temperature (Fig. 2C). The nuclei count gradually decreased with the extension of dissociation time at 37 °C (Fig. 2D), but was not observed in our method (Fig. 2E) and with the intact morphology throughout. Since the release and disappearance of nuclei occur simultaneously during enzyme dissociation, the traditional HED method will force the preferentially obtained nuclei to digest in enzyme solution, or damage the nuclear membrane, affecting nuclear yield, and this method was very sensitive to reaction time, increasing the burden on the experimenter. In addition, the CED method could obtain more than 100,000 nuclei per gram of hippocampal tissue, which was more than 10 times that of commercially nuclear extraction kits based on mechanical methods (Fig. 2F, Fig. S2I). Finally, given the clinical demand for snRNA-seq in a variety of organs, we also dissociated the nuclei of multiple organs (Soybean size) by CED method, including heart, liver, spleen, lung, stomach, intestines, kidney, and pancreas (Fig. S2). We observed strong applicability of this approach to multiple organs, except to the heart and lung. Especially in spleen, intestines and kidney, with the tens of millions of nuclei numbers, and despite their abundance and dense arrangement, they remained independent, intact, and unaggregated (Fig. S2C, F and G). Despite the lower fitness of CED method in the heart and lung, outperformed the mechanical homogenization (Fig. S2H). Moreover, conventional wisdom suggests that nuclei are not freeze-thaw friendly, forced to improve the experimenter’s awareness of time control. Notably, nuclear envelope rupture and aggregation did not occur in nuclei isolated by CED even after one month of dry ice or storage at −80°C. This property breaks the restriction that the nucleus cannot be cryopreserved.

### 2.2 Without damage of RNA molecules in the nucleus from CED method

The morphological of the nucleus ensures the independence of single nucleus data, while the quality of RNA molecules in the nucleus can ensure the high-quality output of snRNA-seq, which is also one of most important factors affecting its application in transcriptome research. PFA fixation of cells induces cross-linking between nucleic acids and proteins, whereas the preparation of FFPE samples requires hours of high temperature wax immersion, both result in RNA damage. How to avoid secondary damage to RNAs during nuclear preparation is crucial for snRNA-seq. We extracted RNA from nuclei to verify the harmlessness of CED method on RNA molecules. We first investigated how to extract RNA molecules from the nuclei of fixed brains. The conventional commercial RNA extraction kits are obviously not suitable for the nuclei of PFA-fixed and FFPE tissues. We combined two lysis systems which suitable for fresh or frozen tissue to extract RNA from cross-linked nuclei. The effects of different heating conditions and proteinase K concentrations on RNA integrity (RIN) and RNA yield were tested using Drop-seq buffer and commercial RNA extraction kits. We found that PFA cross-linking was effectively reversed by incubation at 56 °C for 15 min in standard Drop-seq lysis buffer (Fig. S1B), significantly shortening the heating time compared to the reported [18–20]. PK has been reported to increase RNA yield [18, 19], but our results shown that the PK concentration has little effect on RIN and RNA yield (Fig. S1C). In addition, we performed the same experimental exploration on lysis systems of other high-throughput sequencing platforms, although comparable amount of RNA could also be obtained, RIN values were low (2-4). This means that the standard Drop-seq lysis buffer can be used directly as lysate for FFPE nuclei at 56 °C.

The results showed that the CED method had almost no damage to the RNA molecules compared with the HED method. The RIN values of nuclei were basically consistent with the RIN values of PFA-fixed tissues [PFA(4T) vs PFA-fixed section] (Fig. S1D), but far higher than that of the nuclei prepared by HED method [PFA(37T)], and even the RIN values of FFPE [FFPE(4T)] nuclei was higher than that of PFA(37T) (Fig. 2F). Then, the cDNA libraries were generated from multiple tissues to truly reflect the quality of polyA_RNA. The major peak size of cDNA for both PFA(4T) and FF(0h) was above 1200bp, while around 800bp for PFA(37T), which was even lower than that of FFPE(4T) (Fig. 2H), which again confirmed that CED method was less damaging to RNA molecules in the nucleus than the HED method. Notably, the RNA yield of nuclei isolated by CED method was consistent with that of freshly frozen (FF) samples, but much higher than that of traditional methods (Fig. 2G), which might be a key reason for limiting the application of snRNA-seq for post-fixed samples. We found that free RNA penetrates into the enzyme solution during high-temperature dissociation process, resulting in the reduction of the amount of RNA in the nucleus (Fig. S1E), which we concluded to be a fatal shortcoming of the conventional method. It has been reported that 3× or 5× SSC proved to be a good medium for the prevention of cellular RNA degradation, but we found that that the two types of buffers had almost the same effect on RNA molecules (Fig. S1F). In conclusion, CED method can minimize the damage to the nuclear morphology and RNA molecules of post-fixed brains.

### 2.3 Validation the nuclei quality derived from CED method by snRNA-seq

We employed droplet-based snRNA-seq technology to capture total RNA (M20 Genomics) and poly(A) RNA (10× Genomics). snRNA-seq was performed on mouse hippocampus samples with three treatment conditions: (1) fresh frozen tissue-the nuclei were extracted by mechanical homogenization, capture by poly(T) [FF (10×)] and random primers [FF(M20)]; (2) PFA-fixed tissue - the nuclei were prepared by HED [PFA(37T)] and CED method [and PFA(4T)] for snRandom-seq; (3) The nuclei of FFPE tissue were dissociated by CED method and capture by random primers [FFPE(4T)/FFPE]. With snRNA-seq data of frozen tissue for reference, the influence of high and low temperature dissociation on sequencing data was evaluated to verify the fidelity of nuclei obtained by the CED method, and also to evaluate the applicability of CDE method in FFPE samples. Before microfluidic encapsulation, the nuclei were imaged to confirm single nucleus morphology and counted, and the results returned that the nuclei numbers of all samples were about million level. snRandom-seq necessitate a substantial amount of input material, millions of nuclei fully meet the requirement of nuclear detection rate of 10,000. After barcoding and amplification, the fragment size of the cDNA from FF(M20) main peaked about at 700 bps (Fig. S3A). While the main peak of cDNA from PFA(4T) between 300 and 1000bps, and longer than PFA(37T), which might be stem from RNA degradation (Fig. S3A). In addition, the next-generation sequencing (NGS) library with equal input of cDNA showed the lowest library for PFA(37T) (Fig. S3B). The amount of cDNA and NGS library from PFA(4T) was slightly higher than that in FFPE(4T) and FF(M20), but much higher than PFA(37T), indicating that CED method effectively blocked the leakage of nuclear RNA and almost maintained the true level RNA molecules within the nucleus (Fig. S3B). In fact, we have optimized the HED method to work well for snRNA-seq, which has improved its investigability in transcriptomics, but the results are still unsatisfactory. In short, the nuclei prepared by the CED method from fixed or paraffin-embedded samples are more suitable for the research of snRNA-seq.

### 2.4 Performance of nuclei from FFPE tissues in snRNA-seq

We identified 150,507 high-quality unique nucleus barcodes using the barcode-gene rank plot, with clearly separated of nuclei from background noise, and an average of more than 10,000 nuclei were identified in every sample (Fig. 3A). Gene and UMI count distribution showed that the total UMIs and genes in FF(M20) and PFA(4T) were significantly (*p* < 0.05) higher than in PFA(37T) (Fig. S4A, B), which confirmed that CED method could maximize the retention of RNA molecules. But the total gene numbers detected in all samples were comparable, all above 30,000 (Fig. S4C). In addition, snRNA-seq captured a mean of 1835, 2725, 1347, 2013, 1847 genes and 4189, 11100, 4996, 8759, 7072 UMIs in single nucleus by sequencing average ∼27k, 19k, 13k, 17k, 12k reads per nucleus for FF(10x), FF(M20), PFA(37T), PFA(4T) and FFPE(4T) samples, respectively (Fig. 3B, C). The number of genes and UMIs in FF(M20) was higher than FF(10x), which benefit from the principle of capturing full length and transcripts by random primers. Moreover, the gene and UMI counts per nuclei in PFA(4T) were slightly lower than FF(M20), but about 1.5 to 1.75 times higher than PFA(37T), and even about 2-2.5 times in individual samples, and their numbers in FFPE(4T) were also higher than that in FF(10x) and PFA(37T) (Fig. 3B, C). The saturation analysis showed that FF(M20) and PFA(4T) had the highest sensitivity, followed by FFPE(4T), with 4000 to 5000 detected genes per nuclei, respectively, at a sequencing depth of 30,000 trimmed reads per nuclei, and both exhibited a higher gene detection rate than PFA(37T) (Fig. 3G). We next compared our data with other reported results (Fig. 3D, Fig. S4D). The genes detected per nuclei in snCED-seq datasets of brains comparable with other parenchymatous organs, despite inherently lower RNA abundance in brains (Fig. S4D). The number of genes detected in snRNA-seq reached saturation between 100k and 150k uniquely aligned reads per nuclei (Fig. S4D). Beyond that, lower rates of mitochondrial and ribosomal genes in PFA(4T) and FFPE samples than others, almost 0, indicating that the CED nuclei were pure without cytosolic contamination (Fig. 3E, F). Unlike FF(10x), samples captured by random primers exhibited homogeneous coverage across the body of protein-coding, but with a slight bias toward the 3′-end due to the extra addition of oligo(dT) primer in reverse transcription (Fig. S3E). However, PFA(37T) had more 3′ bias, possibly related to the greater fragmentation of RNA within nuclei (Fig. S3E).

**Fig. 3.**
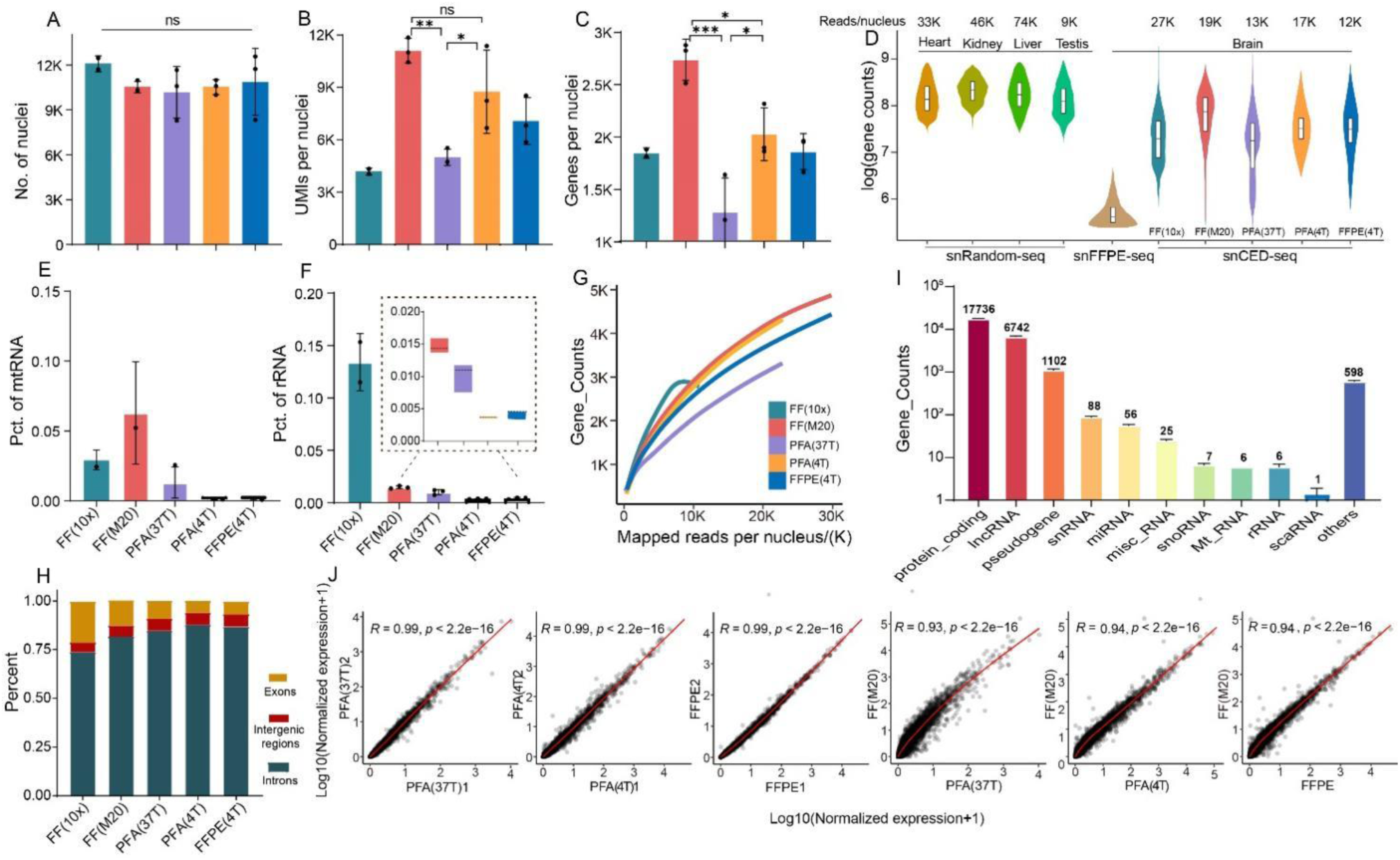
Validation of snCED-seq quality control data. (**A-C**) Number of nuclei (A), UMIs per nuclei (B) and genes (C) per nuclei detected in fresh frozen, PFA-fixed and FFPE samples; (**D**) Gene detection per nuclei comparison of our data (>10,000 nuclei) with mouse tissues (5795 (kidney), 4287 (liver), 6732 (heart) and 3774 (testis) nuelci) by snRandom-seq [10], mouse brain (7031) by snFFPE-seq [9] and breast (5721) by snPATHO-seq [8]; (**E, F**) Percentage of mitochondrial (E) and ribosomal (F) genes; (**G**) Saturation analysis of snCED-seq based on the different samples; (**H**) Percentage of reads mapped to different genomic regions under different conditions; (**I**) Counts of different RNA biotypes detected in FFPE brains. (**J**) The Pearson’s correlation coefficient (R) of the normalized gene expressions between technical replication samples and post-fixed/fresh samples.

In our snRNA sequencing experiment of PFA-fixed and FFPE brains, less than 10 % uniquely aligned reads were mapped to exons and intergenic regions, and with more reads mapped to introns (Fig. 3H). In contrast, frozen samples had a higher proportion of exons (FF(M20): ∼13% and FF(10x): ∼21%) (Fig. 3H). We suspected that the nuclei of frozen samples were prepared by the homogenization method, which was more susceptible to the cytoplasm of pollution. By comparison, higher coverage of intronic regions in the post-fixed groups, especially in PFA(4T) and FFPE(4T) (Fig. 3H), suggesting that our nuclei had little cytosolic contamination, with higher fidelity. The higher proportion of introns might lead to more accurate RNA velocity measurements across differentiation trajectories [21]. Abroad spectrum of RNA biotypes was detected, and protein-coding genes were the most highly detected biotype across all groups, but also other biotypes (Fig. S5A). Unexpectedly, a substantial amount of full transcripts were detected in all group samples, especially in FF(10x) (Fig. S5A), used 10x Chromium Single Cell 3′ Solution, which is consistent with our previous bulk RNA-seq analysis [22]. Contrary to our previous knowledge, we speculated that perhaps there is a wider range of A-capped non-coding RNA molecules within the nucleus. However, at least our data show that extensive and ought to exist non-coding genes can be detected in the FFPE nuclei prepared by CED method (Fig. 3I).

Next, gene expression correlation analysis was performed on our data. To prove the repeatability of our method, duplicate samples were sequenced independently, and a high correlation (Pearson R: 0.99, p<2.2e-16) of gene expression profiles across random batches were seen in PFA(37T), PFA(4T) and FFPE(4T) groups (Fig. 3J), indicating the robustness of nuclei from fixed/FFPE samples. We then analyzed the correlation of gene expression between fixed/FFPE and frozen samples. Consistently, the total RNA profiles of fixed/FFPE and FF(M20) samples displayed a good correlation (PearsonR : > 0.9, p < 2.2e-16), more genes were underexpressed in PFA(37T) group, but not observed in PFA(4T) and FFPE(4T) samples (Fig. 3J). A poor correlation between fixed/FFPE and FF(10x) (PearsonR: ∼0.7, p < 2.2e-16) (Fig. S4F), and higher gene expression in fixed samples, which stem from differences in technique. In addition, the correlation between FFPE(4T) and PFA(4T) was as high as 0.99, reaching the within-group level. Compared with PFA(37T), it was only 0.95, and the gene expression was higher in FFPE(4T) samples (FIG. S4F). These results suggest that nuclei from the CED method behave more similarly to frozen samples.

### 2.5 Cell heterogeneity analysis in PFA-fixed and FFPE tissues

We next compared the cell types identified in all group samples at single-cell resolution. Unsupervised clustering of the above filtered high-quality single brain nucleus profiles, by merging the data of PFA-fixed, FFPE samples and frozen samples. By merging the data of all batch of samples, we obtained a robust cell clustering by UMAP (Uniform Manifold Approximation and Projection), and the low similarity cellular landscapes between FF and fixed/FFPE samples before batch (Fig. S6A). Batch-based processing resulted in integrated UAMP profiles revealed over 21 distinct clusters (Fig. 4A, Fig. S6B). All clusters could be further annotated based on classical known cell-type markers (Fig. 4B), and 11 major cell types were identified with cell-specific genes reliably mapped on the corresponding clusters (Fig. 4B, Fig. S6C). Most of the recommended terms in mouse hippocampus samples were identified, including excitatory neuron (Ex1-8), inhibitory neuron (Inh1-4), Interneuron (Inter_N), astrocytes (AST), Oligodendrocytes (Oligo), Oligodendrocyte pregenitor cells (OPC), Microglial (Micro) and Cajal Retzius cells (CRC) (Fig. 4A). Besides the known cell types, we also respectively annotated choroid plexus cells (CPC) markered by *Prlr*, which are rarely detected in reported data (Fig. 4A). We suspect that our nuclei were more abundant and contained more cell types.

**Fig. 4.**
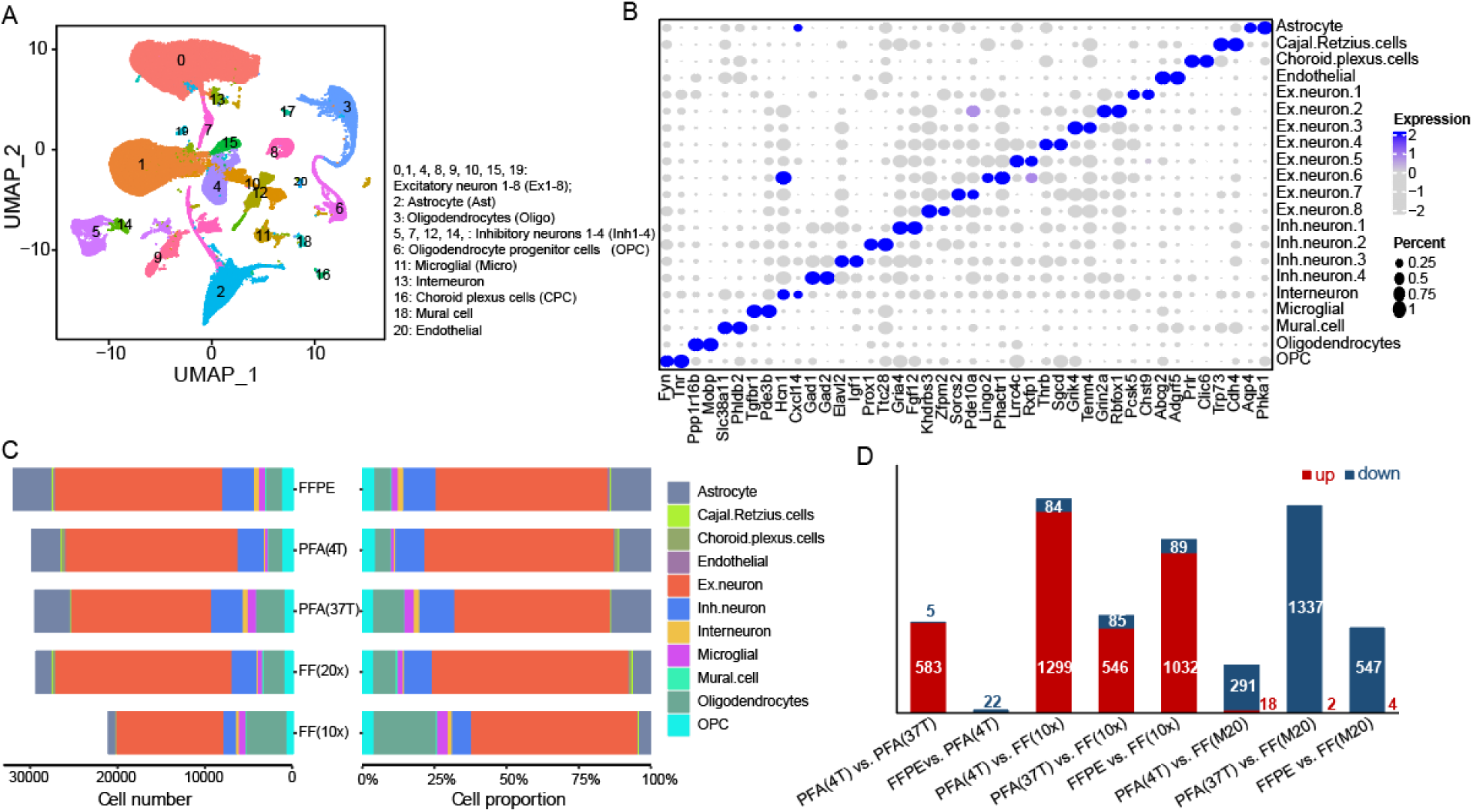
snCED-seq distinguishes major brain-cell types. (**A**) Cell map of mouse hippocampus. UMAP of 150,507 single-nucleus RNA profiles from hippocampi of fresh frozen, PFA-fixed and FFPE samples. colored by cluster; (**B**) Dot plot of the average expressions of top two markers in each of the 21 clusters. The color bars indicate the gene expression level, the bluer the color, the higher the expression level. The bubble diameter indicates the proportion of expression of the gene in that cell cluster, and the larger the diameter, the stronger the specific expression. (**C**) Number of nuclei (Left) and proportion (Right) of annotated cell types of all samples by snCED-seq; (**D**) The number of differentially expressed genes (DEGs) between each comparison group; Red and blue indicate up-regulation and down-regulation, respectively.

Subsequently, we analyzed the proportion of cells types across all groups. As expected, the proportion of cells differed between 3’ and random primer capture techniques, mainly in AST, Oligo, and Endo cells (Fig. 4C, Fig. S6D). However, the similarly cell proportion between frozen and post-fixed samples in our datasets was seen (Fig. 4C). We surmise that we shortened the time of enzymatic dissociation of the samples at high temperatures, thereby retaining most of the cell types in PFA(37T) group. However, the experimentalists are required to be experienced, otherwise resulting in poor batch. Despite this, CPC cells were severely lost in PFA(37T) samples, but a considerable cells were detected in both PFA(4T) and FFPE samples (Fig. S6D), indicating that CED detected more scarce cells than conventional methods. In addition, higher number of cell clusters was obtained at a resolution of 0.1 than other reported results [23–25]. Therefore, we counted the cluster numbers under different resolution and found that absolute advantage in PFA(4T) samples, and even a higher cluster number of FFPE sample than PFA(37T) group (Fig. S6E). But the number of clusters reached a comparable level in all samples when resolution at 1.0 (Fig. S6E). We reconfirmed the above inference that the nucleus prepared by CED method may bring more cellular heterogeneity information and has the potential to recognize more cell types.

### 2.6 snCED-seq revealed cell diversity and heterogeneity in FFPE hippocampal from AD mice

To validate the promise of our nuclei for the research of brain diseases, we applied snCED-seq on the FFPE hippocampus of AD and matched wild type (WT) mice (Fig. 5A) to explore the specific-cell state changes of AD samples. After nuclei with over or under expression of genes were filtered out, snCED-seq identified 62,000 true nuclei in the FFPE brains, and with approximately zero mitochondrial and ribosomal genes in all samples (Fig. S7A). Unsupervised clustering of the single nucleus revealed 19 distinct clusters at resolution of 0.1 (Fig. S7B). The main cell types of AD and WT hippocampus could be identified based on the known cell-type markers, including Ex1-6 (*Hs6st3*, *Pdzrn3*), Inh1-4 (*Gad1*, *Gad2*), AST (*Slc1a2*), Oligo (*Mbp*, *Mobp*), Micro (*Dock2*), OPC (*Vcan*), Endo (*Flt1, Mecom*), Smooth muscle cell (SMC, *Ebf1*). Olfactory ensheathing glia (OEG, *Bnc2*) and CRC (*Cdh4*, *Reln*) with minimal number of cells were also identified (Fig. S7C).

**Fig. 5.**
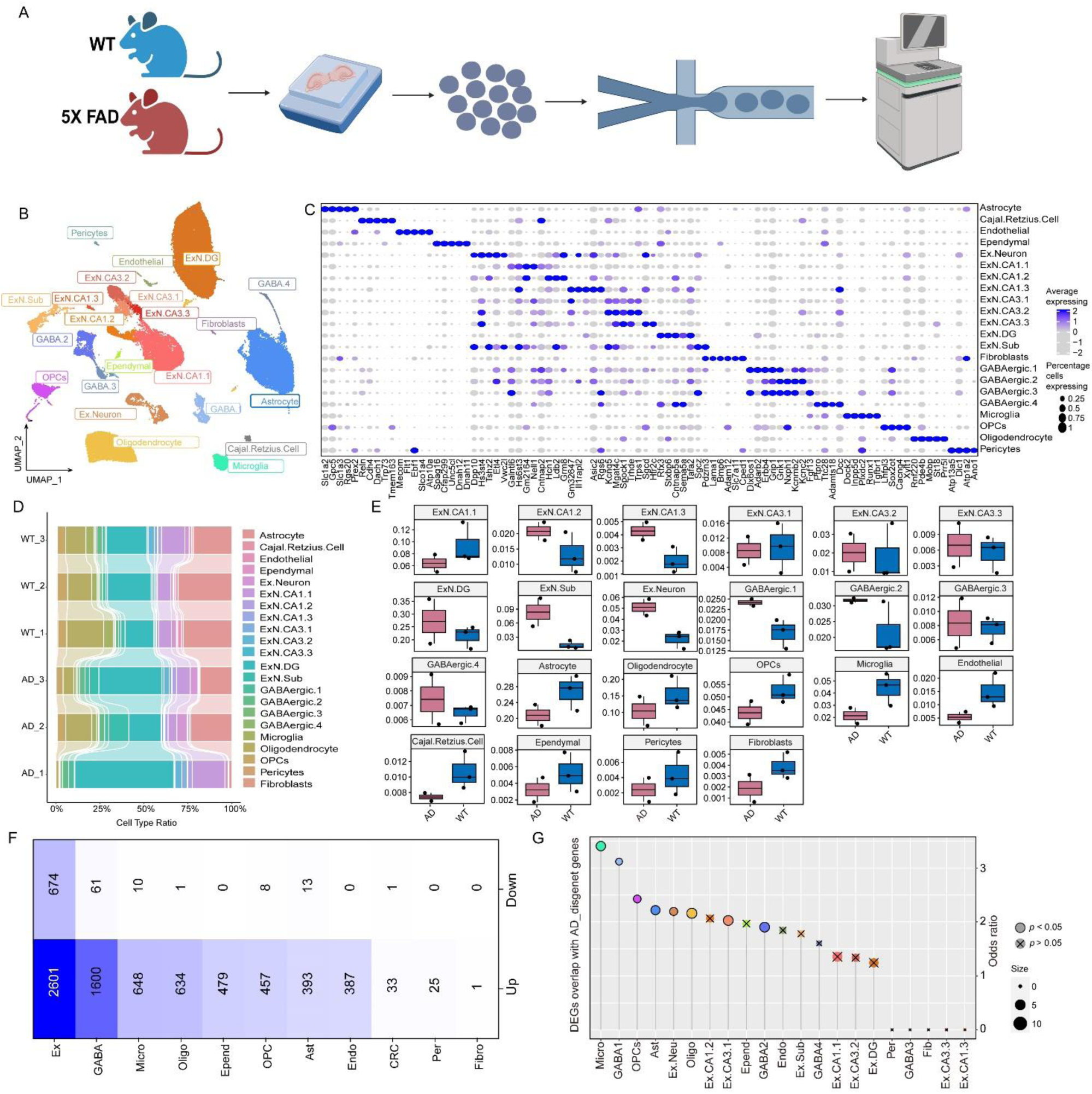
snCED-seq distinguishes major cell types and shows disease-cells in the 5XFAD brains. (**A**) Overview of the experimental strategy. (**B**) Cell map of mouse hippocampus in WT and AD by supervised clustering with reference[25]. UMAP of 62,000 single-nucleus RNA profiles from hippocampi of 5-month-old male mice, three WT and three 5xFAD (AD); colored by cluster. (**C**) Heat map showing expression of specific markers in all cell types, identifying each cluster in B. Expression level (color scale) of marker genes across clusters and the percentage of cells expressing them (dot size). (**D**) The frequency of each cluster in every sample. (**E**) The percentage of cell types in AD and WT. AD1 sample was screened. (**F**) DEG counts for each cell type The intensity of the blue colour was proportional to entry values. (**G**) The odds ratios of DEGs and AD-disease genes in every cluster. The dot size expresses cells association with the AD disease.

Abundant cell types could be identified in our data, but differences in cell proportions compared with previous data from frozen hippocampal of 7-month-old mice [2], such as an increased proportion of most neuronal cells in the AD model, but AD-related cells (Ast, Micro, Oligo, OPC) and vascular-related cells (Endo, SMC) were absent (Fig. S7F). The AST proportion gradually increased with the increase of age of AD [25], and a decrease in the proportion of AST of AD mice has also been reported [23]. We speculate that it might be due to inconsistent methods of nuclear preparation or age of AD mice.

To identify cell types more accurately and precisely, a reference of the published snRNA-seq data of frozen AD hippocampal [25] was used for supervised clustering using Approximate Nearest Neighbors Oh Yeah (Annoy) (Fig. 5B). 90.7 % of the FFPE data and the reference data were predicted to be high predicted scores, and only a minority of the cells had a score less than 0.8 (Fig. S7D, E). However, we could also infer cell types from the distribution of low-predicted cells in the UAMP map, such as Ast and ExN cells (Fig. S8A). The gene and UMI numbers in all cell types from the high predicted score was higher, and the other quality control data was also better (Fig. S8B). In addition, the ExN.IEGs cells from the reference data were successfully detected in our data, but in a small proportion, and overlapped with ExN.CA1.1 cells, so we included them in the ExN.CA1.1 (Fig. S8C). Finally, we determined the atlas of supervised clustering, and identified 22 clusters covering 11 cell types, including 9 Ex cells (ExN, ExN.CA1.1-1.3, ExN.CA3.1-3.3, ExN.DG and ExN.sub), 4 Inh cells (GABAergic.1-4, GABA1-4), and 9 non-neuronal cells (Fig. 5B, 5C). We observed a disproportion of cells in the AD1 that did not conform to conventional wisdom (Fig. 5D), but the proportion of diseased cells before and after removal of the AD1 sample was barely affected (Fig. 5E, Fig. S7G). Overall, the cell types in the reference data were all detectable in our FFPE nuclei, with Ex cells accounted for the largest proportion (52 %), followed by Ast (21 %), Oligo (11 %), GABA (6 %), OPC (4 %) and Micro (3 %), and the other cells accounted for about 1 % respectively (Fig. S8D).

The proportions of cells obtained by both clustering methods were similar (Fig. S7F, S7G). We observed that most cells coincided with unsupervised clustering by marker gene comparison, with ExN.DG corresponding to Ex.neuron6 and GABA4 corresponding to Inh.neuron2 (Fig. 5C, S7C). However, the annotation of some cells changed. For example, we merged Ast1-2 in the reference data into AST, and labeled SMCs and OEGs in unsupervised clustering as pericytes_Per and Fibroblasts_Fib, respectively (Fig. 5C, S8C). OECs are a glial cell between Schwann cells and oligo, which have the functions of neurotrophic, inhibition of gliosis, scar formation and sheath formation, and can provide a suitable microenvironment for axon growth and strong migration characteristics. It has been reported that OECs transplanation reduced amyloid burden in amyloid precursor protein transgenic mouse model [26]. OECs injected into the hippocampus of AD mice can improve the learning and memory ability and increase the activity of mitochondrial cytochrome oxidase in the hippocampal CA1 region, which has an obvious therapeutic effect on AD. This is consistent with our results that OECs in AD undergo loss (Fig. S7F). Notably, CR cells are only present in our data other than reference data (Fig. S7D). The number of CR cells decreases with brain development, and a handful of CR cells can still be detected in the hippocampus of old mice [27]. Since the dominant advantage of the CED strategy to maximize the retention of nuclei of FFPE tissues, relatively few CR cells distributed in the hippocampus were efficiently enriched, and detected by snRNA-seq. We then observed that the proportion of all glial and vascular-associated and other nonneuronal cells were reduced in AD, compared with WT (Fig. 5E). Although the proliferation of Ast and Micro cells is deemed to be the cellular changes of AD disease, but the frozen hippocampal snRNA-seq data reported by Regev, except for Micro and Endo cells, the remaining proportion changes of non-neural cells are consistent with ours [25]. Also, in the cortical data, Ast was in a status of missing, Micro was the only cell type that increased in AD, and the rest were in a stable state. But in the hippocampus, Ast, OPC and vascular cells were reduced in AD, the proportion of Micro increased, and Oligo remained almost unchanged [23]. The similar results further demonstrate the reliability of our data.

In brief, more abundant cell types can be detected in our data, and provide the superior nuclei for omics research in brain diseases. We next used these cells to characterize the heterogeneity of AD- disease cells and the perturbing nature of perturbation of gene expression.

### 2.7 Multidimensional identification of AD disease-specific cells

To reveal AD-associated cells, We compared levels of gene expression in nuclei isolated from AD versus WT individuals by cell type, and identified 8026 unique differentially expressed genes (DEGs) that implicated all major cell types, and 90 % of DEGs were overexpressed genes (Fig. 5F). Neurons showed a strong signature of activation, 79 % of DEGs in Ex (Ex.CA3 predominated) and 96 % in GABA (GABA2 predominated) neurons were overexpression, whereas DEGs of non-neuronal cells were almost 100 % upregulated (log2FC> 0.25, *p* < 0.01) (Fig. S9A). Indicating that most genes in AD nuclei were in an activated state. Both up- and down-regulated DEGs were highly cell type specific, 62% of DEGs in neurons, whereas DEGs in non-neuronal populations were substantially smaller, probably owing to reduced power in lower-abundance cell types [28]. Furthermore, vascular cells (Endo and Epend) also showed no less differential changes than glial cells (Fig. 5F, Fig. S9A). These contrasting observations on the number and dominant directionality of DEGs reveal a heterogeneous response to AD between cell types-a recurrent theme that will be observed throughout the study.

The vast majority of DEGs (50%) were perturbed only in a single cell type, which indicates that these perturbations are strongly cell-type specific (Fig. S9B). But a thimbleful of genes was highly expressed in 82 % cell types, such as *Magi2*, *Cadm2*, *Grm7*, *Adgrl3*, *Ctnna2*, *Ctnnd2*, *Camta1*, *Dgki*, *Drc1*, *Lsamp*, *Mbd5*, *Nrg3*, *Ppfia2* and *Prkce* (Table S2). *Rn18s-rs5* and *Malat1* genes were underexpressed in most cells. Among them, *Magi2* has been reported to be associated with AD phenotypes [190] and is considered a potential “new” candidate locus in the etiology of divergent AD, which was involved in the regulation of protein degradation, apoptosis, neuron loss, and neurodevelopment [191]. We speculate that these genes preferentially undergo perturb changes in expression in AD pathology, which may be therapeutic targets associated with AD disease. Overall, these results of our snRNA-seq for FFPE brains indicate that all major cell types are affected at the transcriptional level by AD pathology. Finally, we evaluated whether AD-associated variants are enriched in genomic regions with genes whose expression pattern is cell-type-specific. Fisher test enrichment scores of each cell type-specific DEGs and AD risk genes were calculated, and AD risk variants were found to be associated with genes from Micro, OPC, Ast and Oligo cells, and were also significantly (*p* < 0.05) enriched in GABA1 and GABA2 (Fig. 5G).

The multi-dimensional analyses results showed that nuclei prepared by CED method could identify disease-specific cell types counterpart to frozen samples by snRNA-seq, and vascular cells, which have been less studied in AD, also surfaced. Next, the traditional AD-associated cells (Micor, Ast, Oligo) were used for further analysis firstly.

### 2.8 Microglial heterogeneity analysis associated with AD-related traits

Using the single-cell resolution feature, we sub-clustered Micro cells of AD and WT mice, and identified four subpopulations (Micro0-3) (Fig. 6A). The micro2 was representative cells only in 5xFAD mouse, but micro0 and micro1 were mainly distributed in WT (Fig. 6B). A scanty of 15 micro3 cells were distributed independently in the UMAP atlas, and with the 1 to 2 ratio in AD and WT (Fig. 6B), indicating that Micro cells are significantly affected by AD pathology and the disease-induced differences result in two nonoverlapping cellular states. We observed that the top 10 up-regulated DEGs in AD were highly expressed only in Micro2 and Micro3, implying specific disease changes for these cells (Fig. 6C). Micro’s DEGs overlapped with the AD disease gene set, and 11 genes were identified (Fig. 6D). These genes were mainly expressed in Micro2 and Micro3, and individual genes were up-regulated in Micro1 (Fig. 6E). Hence, we determined Micro2 is a DAM (diseases-associated microglial, DAM). Although Micro3 cells accounted for less than 1% of the total, disease genes were specifically highly expressed in them, such as the β-amyloid precursor protein related gene (App) (Fig. 6E). We found that the expression of these genes was higher in Micro3 of AD mice (Fig. 6F), implying that five Micro3 cells in AD were also DAM cells, indicating that our nuclei are highly cellular heterogeneous and more suitable for the application of transcriptomics in diseases.

**Fig. 6.**
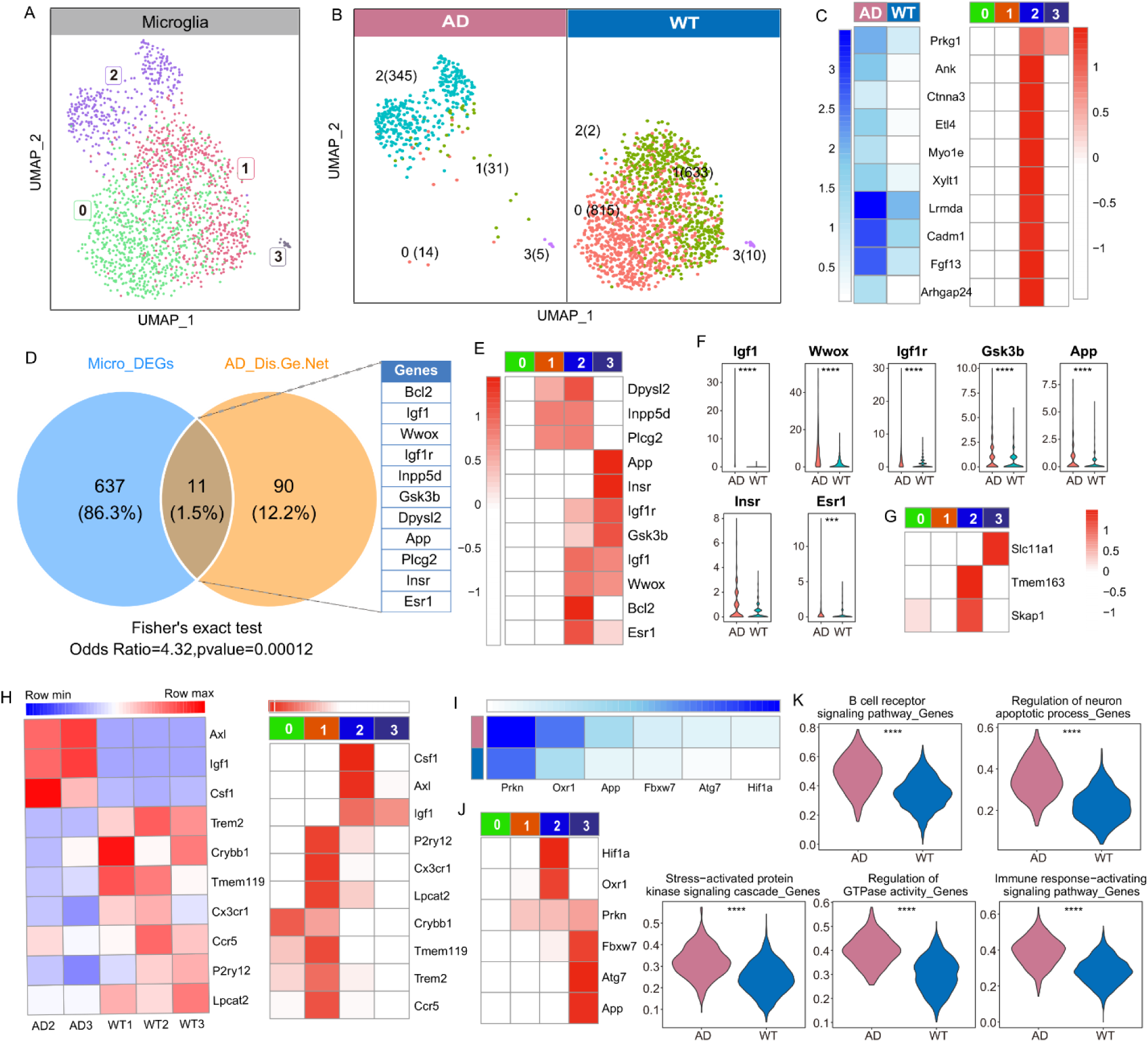
Characterization of the DAMs in AD. (**A**) UMAP plot of re-clustered microglia identifying 4 sub-clusters. (**B**) Cell map of mouse hippocampus in WT (Left) and AD (Right). (**C**) Average scaled expression of the top-10 upregulated disease-specific DEGs in split by sample (Left) and cluster (Right). (**D**) Venn diagram of DEGs in microglia with AD disease gene sets. (**E**) Heat map of intersection genes expressed in Microglia sub-clusters. (**F**) Violin plot of genes in subcluster 3 highly expressed in AD. (**G**) Heat map of amyloid-related genes expression in sub-clusters. (**H**) DAM genes specificity high expression in Micro 2 and Micro 3. (**I-J**) The genes associated with disease-related function or pathway were highly expressed in AD (I) and all clusters (J, K). n = 3 biologically independent mouse brain samples per genotype; Color scheme of heat maps shows row max and row min, which represents relative expression of each gene among AD and all sub-clusters.

To demonstrate the accuracy of the DAM cells we identified, we employed multi-channel data for validation. The expression of DAM genes in human cerebral cortex was first verified in our data (Fig. 6G). The encoded proton divalent cation transporter *Slc11a1*[203], which regulates ion homeostasis and has pleiotropic effects on proinflammatory responses, was expressed only in Micro3. The zinc efflux transporter gene *Tmem163* and immune cell adaptor gene *Skap1* were expressed only in Micro2, which were specific for AD brain (Fig. 6G). SKAP1 is an immune-cell adaptor that couples T-cell receptors to the “inside-out” signaling pathway of LFA-1-mediated T-cell adhesion. Studies have reported that *Skap1*-deficient mice are highly resistant to collagen-induced arthritis, which is a new potential target for therapeutic intervention of autoimmune and inflammatory diseases [29]. Thus, the high expression of *Skap1* in Micro2, lose its anti-inflammatory resistance, which is promising to be a new checkpoint for studying the mechanism of AD disease. Then, DAM genes in cortical Micro of 7-month-old 5XFAD mice were also used to verify the accuracy of our DAM identification [23]. The results showed that *Axl*, *Lgf1*, and *Csf1*, which were upregulated in AD cortex, were all highly expressed in AD hippocampus compared with WT, but the homeostatic genes, such as *P2ry12*, *Crybb1*, *Tmem119* and *Cx3cr1*, were under expressed in AD (Fig. 6H, Left). However, *Trem2* was under-expression in AD of our data, whereas *Csf1*, a gene reported *Trem2*-dependent upregulation, does not appear to be affected by *Trem2* deficiency in the AD hippocampus. Similar to the above result, the reported DAM genes were mainly highly expressed in Micro2 and Micro3, while most of the down-regulated genes were more prominently expressed in Micro1 (Fig. 6H, Right). The expression profiles of reported disease-pathway genes associated with AD in microglia subclusters were also analyzed. The antioxidant defense system is essential for cell survival in the central nervous system, and oxidative stress dysfunction is associated with neurodegenerative diseases [30]. Therefore, we first analyzed the genes involved in the regulatory pathway of oxidative stress-induced neuronal death. We found that these genes were highly expressed in AD mice (Fig. 6I), such as the amyloid gene (*App*), the ubiquitin protein ligase gene (*Prkn*), and the oxidation resistance 1 gene (*Oxr1*). Analogous results were presented again, that all the genes related to this pathway were significantly (*p* < 0.05) highly expressed in Micro2 and Micro3 (Fig. 6J). In addition, genes involved in the B-cell receptor signaling pathway, regulation of neuronal apoptotic processes, stress-activated protein kinase signaling cascade, regulation of GTPase activity, and immune response-activation signaling pathways were all significantly overexpressed in the AD hippocampus (Fig. 6K).

Our snRNA-seq data from FFPE hippocampal nuclei prepared by CED method can identified two DAM cells, in which Micro2 was the proliferating DAM in AD, and DAM-signature gene expression was independent of *Trem2* expression. Moreover, Micro3 is also affected by AD development and exists in AD independent of Micro2. In addition, the combined analysis of multiple data indicated a strong robustness of our nuclei, which will be verified several times in subsequent analyses.

### 2.9 Similar disease-related transcriptional changes occur in astrocytes and oligodendrocytes

Next, we identified five unique AST transcriptomically defined clusters characterized by high expression of *Luzp2* (Ast0), *Rgs6* (Ast1), *Kcnip4* (Ast2), *Cdh4* (Ast3), and *Rnf213* (Ast4) (Fig. S10A, S10B). The DEGs of AST subclusters between AD and WT were evaluated, and most DEGs were upregulated in AD (Fig. S10C). We identified DEGs that were unique to single or combinations of AST subclusters (Fig. S11A) and evaluated these gene sets by GO analysis (Fig. S11B). We observed that the specific DEG numbers (Fig. S12A) and their enriched GO terms (Fig. S11B) were largest in AST2, and the greatest change (log2FC) in specific expression (Fig. S10C). In addition, when comparing the top-10 up-/downregulated DEGs by cluster and disease state, few conserved/common transcriptomic changes were found across all AST subpopulations but instead found highly cluster-specific transcriptomic changes based on disease state (Fig. S10D). The perturbation of gene expression changes in Ast2 and Ast4 was the most prominent (Fig. S10D), and was provisionally defined as DAA (Disease-associated astrocytes). The DEGs that were significantly highly expressed were exactly the marker genes of Ast2 (Fig. S10B-D, Fig. S11C). Despite the overall absence of Ast cells in AD, Ast2 was highly enriched in 5xFAD mice, and Ast4 also exhibited slight cellular proliferation (Fig. S10E), consistent with the pathological features of astrogliosis in AD. DAA genes, *Kcnip4*, *Erc2*, *Nrg3*, *Nrxn3*, and *Csmd1* were notably highly expressed in Ast2 ofAD mice compared with other subclusters (Fig. S10G). We suspected that these DAA genes were primed activated in the hippocampus, and were preferentially activated during disease induction to dominate cell state changes.

DAA cells of Gfap-high state in the report [25], its upregulated genes were highly and unique expression in Ast4, such as *Gfap*, *Aqp4*, *C4b*, and the gene encoding a serine protease inhibitor linked to increased amyloid accumulation (*Serpina3n*) and encoding a lysosomal cysteine protease involved in proteolytic processing of amyloid precursor protein, *Ctsb* (Cathepsin B)(Fig. S10F). Morover, a set of genes including those involved in endocytosis (*Vim*), complement cascade (*Osmr*) and senescence (*Ggta1*) were also overexpression in AST4, confirming our AST4 as a DAA (Fig. S10F). *Gsk3b* (glycogen synthase kinase 3β gene), *Psen1* (presenile factor gene), *Bdnf* (brain-derived neurotrophic factor), and AD risk gene *Sorl1* (encoding endosomal recycling receptor gene) and *App*, associated with AD pathological pathways were also highly expressed in our two DAA cells (Fig. S10H-I). Then, we examined the expression levels of RNA signatures from bulk datasets, only ischemic related genes (*Mcao*) and inflammation related genes (*Lps*) were overexpressed in Ast2, but downregulated in Ast4 (Fig. S11D).

Following the approach of AST, we also characterized the Oligo subpopulations (Fig. S13A). Six Oligo subclusters were identified, and Oligo2 was characterized by *Kcnip4*, *Nrg3*, *Csmd1* and *Grin2a*, which was consistent with Ast2 (Fig. S13B). Moreover, the top up-regulated genes in AD were similarly distributed in the Oligo2 (Fig. S13D), and its unique DEG and GO term numbers were most than other sub-cells (Fig. S13C, S14A), while the down-regulated DEGs were mainly derived from Oligo5 (Fig. S14C). Unlike Miro and AST, comparable cell proportions of Oligo subclusters between AD and WT (Fig. S13E), which was consistent with conventional cognition. We also observed that AD-related genes were overexpressed in Oligo2 (Fig. S13G, H). Strikingly, marker genes of Oligo5, such as *Gpc5*, *Ntm*, *Rora* and *Nrxn1*, were also highly expressed in Ast4, which was specifically expressed by pathologically (Fig. S13I). Among them, *Gpc5* (Glypican 5) was the susceptibility gene for inflammatory demyelinating diseases [31]. *Ntm* was involved in the negative regulation of neuronal projection development and acts upstream or within cell adhesion. Downregulation of *Rora* inhibits glioma proliferation through NF-κB signaling pathway [32], and its regulatory effect was lost when Ast4 and Oligo5 are upregulated. *Nrxn1* (Neurophin 1) was a cell adhesion molecule that plays a key role in establishing and maintaining synaptic connections, and its abnormal expression has been implicated in schizophrenia [33]. Next, we analyzed the expression of marker genes from AD-pathology-associated Oligo [28] in Oligo0-5, and most of genes were more significantly expressed in the Oligo2 and Oligo5 (Fig. S13J). In particularly, *Qdpr*, *Nlgn1*, *Lama2* and *Fchsd2*, closely related to AD-pathology genes reported, were highly expressed in Oligo5 (Fig. S13J).

Finally, to further confirm the accuracy of DAA and DAO (Disease-associated Oligodendrocytes) identification, we collected functional terms associated with AD pathology of previously reported, and analyzed the enrichment of these disease functions in Ast and Oligo subpopulations (Fig. S12). The results showed that almost all functions were enriched in the Ast2, including lipids, glial cell regeneration, endocytosis, NFκB, endothelial cell differentiation and cognition (Fig. S12A). Ast4 cells, however, were enriched with relatively independent functional sets, including functions in the regulation and regulation of growth, response to oxygen levels, and autophagy (Fig. S12A). Moreover, GO terms enriched in Ast2 were also enriched in Oligo2 with stronger significance and enrichment index (Fig. S12B). And the DEGs of Oligo2 were also enriched in autophagy, apoptosis, mRNA regulation and myelination (Fig. S12B).

In a nutshell, the characteristics of DAA and DAO were similar in our snRNA-seq data of FFPE samples. Oligo2 and Ast2 had the same specific expression genes and transcription differences, while Ast4 and Oligo5 have similar transcription characteristics. We conjectured that a group of disease-susceptible gene sets caused similar transcriptional changes in different cell types, which in turn affected the occurrence and progression of AD.

### 2.10 Integration of astrocytes and oligodendrocytes from multiple datasets

Given the abundance of high-quality, well-powered AD sample AST and Oligo snRNA-seq datasets in the literature, we next sought to determine whether we could resolve the same transcriptomic differences previously reported, and in turn demonstrate the availability of our nucleus. We evaluated AST and Oligo subtypes in each individual dataset and compared them with ours. We compared five AST clusters (G0-G4) in the Grubman dataset [34], four AST clusters (M0–M3) in the Mathys dataset [28], and 7 AST clusters (Z0–Z9) in the Zhou dataset [23], and 9 AST clusters (L0–L8) in the Liddelow dataset [35] were integrated with our AST0-4 for analysis, separately (Fig. S11E). Similar analysis was performed on Oligo subtypes (Fig. S14D). Using our AST and Oligo subpopulation profiles as a reference, we identified sub-cells that were also recognizable in the individual datasets. Although a complete one-to-one correspondence was not possible, we still observed that AST and Oligo subtypes were analyzed in the individual data, and disease-associated cells (Ast2, Ast4, Oligo2 and Oligo5) were clearly identified in all datasets, especially in Mathys and Liddelow and Multi-datasets. For example, AST2 was highly correlated with G0, G3, M3, M4, L3-6 cells, while AST4 was more correlated with G1, G4, M0, L7, L8 (Fig. S11E). In contrast, AST0, Oligo0 and Oligo1 showed poor agreement in these datasets. In conclusion, the results of multi-channel data integration analysis of our AST and Oligo subclusters confirmed the aforementioned argument that the diversity of distribution detected in multiple frozen samples could be detected in our data, again demonstrating that our nucleus has cellular diversity, which lays the foundation for the study of disease heterogeneity.

### 2.11 Transcriptional similarities in different disease-specific cell types

Although there are minimal transcriptional changes in neurons and other cells in AD cortex [23], and a recent work also focused only on astrocytes and oligodendrocytes [36]. But the reported data of AD shows that all major cell types are affected by AD pathology at the transcriptional level [28], which was consistent with our results. In our snRNA-seq data of FFPE tissues, the Micro, GABA1, OPCs, AST, Ex.Neu, Oligo, Ex.CA3.1, GABA.2 cells were more perturbed by AD (Fig. S9A). Moreover, more than 400 up-regulated DEGs were being in two vascular related cells, Epend and Endo, which even exceeded AST (Fig. S9A). To test the previous conjecture that there is a disease susceptibility gene-set with consistent transcriptional differences in different cell types. We performed differential analysis of gene expression for all cell types perturbed by AD to explore the transcriptional similarities of disease-related cell types. (Fig. S15A).

We observed that only the DEGs of Micro were cell-specific, while the most significant DEGs of the remaining glial cells and vascular cells were highly heterogeneous, and the top DEGs of neuronal cells also overlapped strongly (Fig. S15B). Since the Log2FC of two vascular cells were too large to annihilating the information of the other cells, we present them independently (Fig. S15C). The positional candidate or therapeutic marker genes, include the immune-related hub genes (*Fgf13* [37] and *Etl4* [38]), the anti-inflammatory gene (*Myo1e*)[39], the multichannel transmembrane tonic transporter gene (*Ank*), and the cadherin-related protein gene (*Ctnna3*)[40] showed the greatest transcriptional changes only in micro cells (Fig. S15B). However, *Kcnip4*, *Grin2a*, and *Lrp1b* were among the most differentially transcribed in other nonneuronal cells. The gene encoding Kv channel interacting protein 4 (*Kcnip4*) was a candidate gene for attention deficit hyperactivity disorder [41]. The inability of the *Kcnip4* isoform to interact with the secretase complex leads to increased secretion of beta-amyloid enriched in the more toxic Aβ-42 species [42]. And it also has been reported that *Kcnip4* interacts with presenilin, and the presenilin gene is associated with early-onset familial AD [43]. Sleep deprivation (SD) could increases the risk of AD, and N-methyl-D-aspartate receptors (NMDAR) is an important cognitive regulator. Specific knockdown of hippocampal astrocytic *Grin2a* (the gene encoding the NMDAR subunit GluN2A) aggravated SD-induced cognitive decline, elevated Aβ, and attenuated the SD-induced increase in autophagy flux [44]. Most of these conclusions were based on the results of immunofluorescence staining, while our snRNA-seq data showed exactly the opposite, the *Grin2a* gene was not only highly expressed in Ast, but also positively expressed in most of the cell types associated with AD (Fig. S15B). The low-density lipoprotein receptor-associated protein 1B (LRP1B) can interact with APP and regulate its processing to Aβ [45]. In summary, the transcriptional profiling of all cell types closely associated with AD reconfirmed our previous hypothesis that a single disease-susceptible gene set causes similar transcriptional changes in different cell types.

## 3 Discussion

In this study, we developed a strategy for high-quality nuclear preparation from FFPE tissues by enzymatic dissociation of archived sample at low temperature without any tedious filtration step, which therefore provides a critical advance to profile single nuclei transcriptome from low-quality biological samples of PFA-fixed or FFPE tissues of. Meanwhile, we performed snRNA-seq on frozen, PFA-fixed and FFPE brains using 10× Genome and snRandom-seq technologies, and performed head-to-head comparison. To prove our method, we performed validation and obtained promising results. snRNA-seq was performed on frozen samples using 10× Genome and snRandom-seq technologies to eliminate platform differences, and the data of frozen samples was used as the gold standard for reference. We used snRandom-seq to perform snRNA-seq on PFA-fixed samples to compare the nucleus preparation strategies, and explored the applicability of snRNA-seq on FFPE samples as well as its application performance in brain diseases. The CED method and snCED-seq represents a significant advance in single-nuclei sequencing, enabling researchers to retrospectively select samples from a large paraffin sample bank, and facilitating mechanistic studies of brain disease samples that are difficult to obtain clinically.

Molecular biological application of FFPE tissues has always been challenging due to the chemical cross-linked and low-quality RNA. Although researchers have gradually become aware of the potential for obtaining expression profiles of individual cells or nuclei FFPE tissues, the approaches of extracting or isolating high-quality nuclei remains challenging. The acquisition of nuclei is one of the important conditions for snRNA-seq of FFPE samples, and its quality directly determines transcriptome analysis. The preparation methods of FFPE sample nucleus are longstanding, and are mainly divided into two categories, hyperthermal enzymatic dissociation strategies and mechanical extraction strategies. Enzymatically obtained nuclei are unhea1rd of in transcriptomics studies. In fact, prolonged high temperature treatment resulted in secondary RNA degradation of FFPE samples, and prolonged exposure of nuclei to the enzyme buffer may increase the permeability of the nuclear membrane, leading to RNA molecules leakage and adversely affecting snRNA-seq experiments performed in droplets. The mechanical homogenization strategy was less damaging to RNA molecules within nucleus. However, tissue homogenization for fixed and FFPE tissues, becomes increasingly challenging due to molecular cross-linking within the nuclei. Firstly, effectively removing excessive tissue debris poses difficulties and leads to severe contamination of the snRNA-seq data [9]; Moreover, the high proportion of rRNA requires additional removal processes when employing total RNA protocols [21, 46]. In our early experiments, the large amount of tissue debris interfered with accurate identification of nuclei due to the molecular cross-linking introduced by formaldehyde fixation, which requires a complicated debris removal process, thus affecting the yield of nuclei and losing smaller nuclei. Currently, the method employed in high-throughput snRNA-seq platforms rely on a combination of enzyme dissociation and homogenate [10]. Despite the optimization of nuclear suspension and RNA quality within the nucleus, their own shortcomings have not been dismissed. Furthermore, all current methods for preparing nuclei from FFPE samples focus on tissue sections (5-100 μm), while disease research often involves the tissue blocks.

Due to the characteristics of dissociation and digestion of nuclei in the preparation process of enzyme dissociation, the traditional high-temperature method makes the nuclei prepared first be digested in the dissociation solution, or the nuclear membrane is damaged, which affects the nuclear yield and is very sensitive to the reaction time, increasing the burden on the experimenter. Our nuclei were obtained by enzymatic hydrolysis of molecularly cross-linked tissues with a single step at low temperature, and without ultracentrifugation through a sucrose cushion and any filtration procedures, thereby maximizing product retention and nucleation rates. Taking a pair of mouse hippocampus as an example, the number of nuclei prepared by CED method was about 10 times that of the traditional method, and CED method can better enrich the small diameter nuclei missed by the traditional method. Most importantly, our CED method can effectively protect the nuclear membrane and maximally retains the nuclear molecules, providing high-fidelity nucleus for snRNA-seq research. For the latest snRNA-seq technology based on random primer capture [10] or gene probe capture [8], it is necessary to input nearly one million nuclei on the premise of ensuring the output of about 10,000 nuclei. Our CED method effectively avoids the current two major nucleus preparation strategies, and can export the nucleus stably without introducing more impurities and damaging the nuclear membrane. The nuclei prepared by our CED method could be successfully preserved or transported on dry ice, which we speculated might be due to the fact that the permeability of the nuclear membrane was not damaged. In addition, our CED method has good applicability to a variety of organs, such as brain, liver, kidney, pancreas, spleen tissues, but slightly poor compatibility with heart and lung, although the yield of nuclei was still higher than that of the mechanical method. Heart has a complex cellular composition, mainly including myocardial tissue, nerve tissue, Purkinje fiber, connective tissue, epithelial tissue, etc. Similarly, the main connective component of the lung is composed of connective tissue, which is rich in collagen fibers, elastic fibers, reticular fibers. Connective tissue and cell relatively dense structure greatly increases the difficulty of the heart and lung tissue, and the choice of operating conditions and enzymes need further adjustment.

The excellent performance of the CED method was maintained for both PFA-fixed and FFPE tissues in our benchmarking effort. Compared with the HED high-throughput snRNA-Seq database of the PFA-fixed samples, snCED-seq outperforms well in various perspectives, supported by the genes and transcripts per nuclei, percent of mitochondrial and ribosomal genes, gene detection sensitivity, gene expression correlation with frozen samples, especially in gene expression richness. High-quality and high-sensitivity snRNA-seq data from post-fixed (PFA-fixed and FFPE) specimens by snCED-seq allows to identify rare cell populations. We further provide a detailed map of cell-type-specific of FFPE samples from AD and WT mice, which highlights the predominance of gene expression richness in our nuclei. Multiple disease-related subpopulations have been successfully identified, and the DAM has transcriptional independence, while the transcriptional similarity between DAA and DAO subpopulations. There is even a population of genes (*Kcnip4*, *Grin2a*, *Lrp1b*, ect.) that are in the waiting state of activation, priority in different cells by the interference of the disease. In short, nuclei from CED method have excellent in revealing cellular heterogeneity, which contributes to the precision diagnosis and treatment to human disease.

Overall, we have proposed a novel method for the preparation of high-fidelity nuclei from post-fixed samples, which outperforms the traditional method in various aspects, and has good compatibility with a variety of FFPE organs. The application of FFPE samples in AD was also investigated, and found that our nuclei have great potential for uncovering disease cellular heterogeneity. The simple experimental protocols and comprehensive transcriptomic information from the FFPE tissues described in this study are expected to enable snCED-seq to large-scale applications in basic and clinical researches in the future. Our nuclear preparation strategy lays the foundation for revealing the transcriptomics and even multi-omics information of FFPE samples.

## 4 Methods

### 4.1 Ethical statement

The study was approved by the animal ethical and welfare committee of Zhongda Hospital Southeast University (approval numbers: 20200104005). All procedures were conducted following the guidelines of the animal ethical and welfare committee of SEU. All applicable institutional and/or national guidelines for the care and use of animals were followed.

### 4.2 Experimental model

Male wildtype (WT) C57BL6/J mice (8 weeks of age) were ordered from Qinglongshan Animal Farm, Nanjing, China. AD and their control mice were purchased from Jiangsu Huachuang sinoPharmaTechCo., Ltd, Taizhou, China. Five-month-old heterozygous 5xFAD transgenic mice (on a C57/BL6 background) co-overexpress mutant forms of human amyloid precursor protein associated with familial AD, the Swedish mutation (K670N/M671L), the Florida mutation (I716V), the London mutation (V717I) and carry two FAD mutations (M146L and L286V) people PSEN1. The expression of both transgenes is regulated by the mouse neurospecific regulatory element *Thy1* promoter to drive transgene overexpression in the brain. Throughout the study, all mice in each experiment were nontransgenic littermates from the same mouse colony.

All the mice were single-housed under standard laboratory conditions, including a 12h light/darkcycle, temperatures of 25 °C with 40 % humidity, with free access to mouse diet and water. The animals were anesthetized with 500 mg/kg tribromoethanol (Sigma, Saint Louis, MO, USA) and were killed by cervical dislocation. After the animals were sacrificed, hippocampi of brain were isolated. Fresh frozen (FF) tissues were obtained by quickly frozen in liquid nitrogen; PFA-fixed (PFA) tissues were prepared by adding PFA (4 %) to the hippocampus and fixed for 20 h at 4 ℃; FFPE samples were prepared by dehydration the fixed hippocampus twice in 70 %, 90 % and 100 % ethanol, respectively, and then clearing with xylene solution for 15 min, twice, followed by paraffin embedding for 2 h (62 °C). Frozen samples and PFA-fixed samples were stored in a −80 °C, and FFPE samples were stored at 4 °C.

### 4.3 Single nuclei isolation from PFA-fixed and FFPE tissues

For frozen samples: the nuclei prepared by the homogenization method by Singleron Biological Tech Co. for 10x Genomics and M20 Genomics for snRandom-seq.

For PFA fixed samples: the tissue was washed three times with 1 mL PBS (1×, pH = 7.4) and cut into 1 mm^3^ pieces in a 2 mL enzyme-free centrifuge tube, adding 1 mL dissociation buffer (1.5 mg/mL Protease K, TE buffer, pH = 8), 0.5 % Sarkosyl), and shaking at low temperature overnight. The supernatant was centrifuged in a 1.5 mL tube, centrifuged at 10000 rpm for 10 min at 4 °C, and discarded supernatant. Then washed nuclei with 1 mL pre-cold PBS (1×, pH = 7.4) twice, centrifuged at 10000 rpm for 10 min at 4 °C. Finally, the nuclei were resuspended in 200 uL of nuclear store buffer (1× PBS, 0.2 U/mL RNase Inhibitor) and stored at −80 °C.

For FFPE samples: the intact hippocampal tissue was trimmed out of the FFPE blocks with a sterilized scalpel and placed in a 2 mL tube, and washed thrice with 1.5 mL Xylene for 2 h at 4 °C to remove the paraffin. The samples were gently redehydrated by immersing the samples in a graded series of ethanol solutions, starting with pure 100 % ethanol and ending with 50 % ethanol (100% × 2, 95 %, 70 %, 50 % × 1) for 1 h, then washed twice with pre-cold water. The steps of nuclei prepared was same as PFA tissue. An aliquot of nuclei was stained with DAPI (4′,6-diamidino-2-phenylindole) staining solution, loaded on a hemocytometer and observed under an inverted fluorescence microscope. Eligible nuclei were stored on dry ice and sent to M20 Genomics.

### 4.4 Library Construction and Sequencing

For frozen samples: Isolated nuclei were subjected to droplet-based 3′ end massively parallelng using Chromium Single Cell 3′ Reagent Kits per the manufacturer’s instructions (10x Genomics).

For PFA-fixed and FFPE samples: In the single-cell transcriptome sequencing experiments of this study, we utilized the VITAcruizer single-cell preparation instrument DP400 (Cat #E20000131, M20 Genomics) to achieve droplet generation, single-cell partition and encapsulation, and nucleic acid capture. The VITApilote high-throughput FFPE single-cell transcriptome kit (Cat #R20121124, M20 Genomics) was employed for pre-library sample processing, single-cell library construction, and purification. Experimental procedures were conducted following the perspecitve kit and instrument manuals The main workflow is outlined below.

Nuclei were removed from dry ice and thawed at 4 °C, and the qualified single nuclei were subjected to snRNA-seq processing according to a previously published snRandom-seq protocol [10]. Random primers were then added for the reverse transcription of total RNA inside the cell nuclei. Subsequently, the generated cDNA fragments were ligated to adaptors inside the cell nuclei. The reverse-transcribed single-nucleus suspension and reagents, along with barcode beads containing cell barcodes and UMIs, were mixed. The mixture underwent encapsulation, capturing and barcoding using the VITAcruizer DP400 instrument. The resulting product underwent extension, resulting in barcoded cDNA strands. Following that, PCR amplification was conducted using the cDNA as the template. The purified products after cDNA amplification were then used to constructed a standard next-generation sequencing library. The constructed single-cell library contained P5 and P7 adapters and was sequenced on a Novaseq 6000 sequencer (Illumina) with 150 bp paired-end reads.

### 4.5 Data analysis for C57BL6/J mice Preprocessing of snRNA-seq data

*Mus_musculus*. GRCm39.109 reference genome was downloaded from ensemble database. Then we used STARsolo module in STAR (2.7.10a) with default parameters to generate the gene expression matrix and filter the valid nuclei. The Seurat v4.2 was applied for the major downstream analysis. Before we started downstream analysis, there are some filtering metrics to guarantee the reliability of each data. detected in fewer than 3 cells were filtered to avoid cellular stochastic events. We deleted mitochondrial genes after the quality control, the left genes used for downstream analysis. For the cell part, we set different filter standards for each dataset according to the UMI and gene numbers distribution to filter low quality cells. Finally, we got 23248 genes and 142661 cells as the expression matrix to do downstream analysis in the method comparison part.

#### Clustering and Cell Annotation

After quality control, unsupervised clustering was performed using Seurat v4.2. A series of preprocessing procedures including normalization, variance stabilization and scaling data, were performed in an R function ‘SCTransform’based on regularized negative binomial regression. Then, we selected 2000 highly variable genes to integrate all sequencing libraries using ‘FindIntegrationAnchors’ and ‘IntegrateData’ functions, followed by the regression of technical noise. Principal component analysis (PCA) was performed using integrated output matrix, and the reasonable principal component (PC) numbers was chosen using the ‘JackStraw’ function. And we chose the top 30 significant PCs for downstream cluster identification and visualization. Clusters were defined based on ‘FindClusters’ function with resolution from 0.1 to 1 with 0.1 as seperation. Uniform Manifold Approximation and Projection (UMAP) was used for the final dimension reduction and visualization. Based on the cluster results with resolution equal to 0.2, we next used ‘FindAllMarkers’ function with MAST algorithm. We ranked the marker genes according to the p-value and log2 fold change (log2 FC) within each cluster and searched top genes in Cell Marker database [47] and Panglao DB [48] databases to annotate cell types of clusters.

#### Differential expression analysis

Within each cluster, we calculated differentially expressed genes (DEGs) between 2 different conditions by using ‘FindMarkers’ function. we used ‘MAST’ setting as well and the Benjamini– Hochberg procedure to adjust p value. Then we set threshold q_adjust < 0.05, absolute value of log2 FC > 0 to filter DEGs. The DEGs functional enrichment analysis based on Gene Ontology (GO) and Kyoto Encyclopedia of Genes and Genomes (KEGG) was applied by an R package ClusterProfile v4.10.0 using a hypergeometric test and corrected for multiple hypothesis by FDR.

### 4.5 Data analysis for AD and WT mice Preprocessing of snRNA-seq data

*Mus_musculus*. GRCm39.109 reference genome was downloaded from ensemble database. Then we used STARsolo module in STAR (2.7.10a) with default parameters to generate the gene expression matrix and filter the valid nuclei. The Seurat v4.4.0 was applied for the major downstream analysis. Before starting the downstream analysis, we used four filtering metrics to ensure the reliability of the data. (1) The detected genes were discarded to less than 3 cells to avoid cell random events; (2) Remove nuclei with mitochondrial gene expression percentage >10% to exclude apoptotic cells; (3) Remove UMI >30000 cells; (4) Remove cells outside the range of 300 to 5,000 genes. After filtering cells and genes based on the above metrics, we further use Doublet Finder V2.0 with default parameters to predict and remove potential doublet in each sample. Only cells that have passed a rigorous multi-step quality control regimen are considered for downstream analysis. Thus, 23,248 genes and 63,789 nuclei were retained in AD part.

#### Clustering and Cell Annotation

After quality control, we aligned the data from different batches using the SCTransform [49] integration workflow in Seurat with default settings. We identified high-resolution clusters (resolution = 0.1) using the Seurat functions FindNeighbors and FindClusters (Leiden clustering algorithm) based on the first 30 principal components. To annotate cell types within this dataset, we employed two distinct approaches: (1) manual annotation using previously published databases Cellmarker 2.0 and and Panglao DB databases, and (2) projecting annotations onto the cells analyzed in this study by integrating the clustering results with the dataset from Habib et al. [25] using the Seurat functions FindTransferAnchors and TransferData. Combining these methods yielded detailed and reliable cell cluster information. We also excluded certain less reliable cell types (e.g., Ex.neuron2 and Ex.IEG). During the subsequent cell proportion analysis, we observed a significant imbalance in the cell proportions of the AD1 sample, with neurons comprising the majority and non-neuronal cells only accounting for 4 %. Thus, we deemed this sample unreliable and excluded it from further analysis. Ultimately, we retained 23,248 genes and 52,569 cells.

For the subcluster analysis of astrocytes, oligodendrocytes, and microglia, we similarly utilized the Seurat functions FindNeighbors and FindClusters on the top 30 principal components, setting the resolution to 0.1.

#### Differential expression analysis

Within each cluster, we used the “FindMarkers” function in the Seurat package to detect DEGs between AD and WT conditions. We applied the “MAST” setting and controlled the false discovery rate (FDR) using the Benjamini-Hochberg procedure. We set thresholds of |avg_log2FC| > 0.25 and p_val_adj < 0.05 to filter DEGs, identifying both upregulated and downregulated genes in AD relative to WT for each cluster.

#### Enrichment analysis

The functional enrichment analysis of Differentially Expressed Genes (DEGs) based on GO biological processes and the KEGG was conducted using the R package ClusterProfile v4.11.1. The analysis employed a hypergeometric test and corrected for multiple hypotheses using the False Discovery Rate (FDR). For enrichment between gene sets, the testGeneOverlap function from the R package GeneOverlap v1.34.0 was utilized to perform Fisher’s exact test, identifying overlaps among different gene sets.

#### Comparison with external data sets

To compare our data with external datasets, we collected single-cell data from studies by Grubman, Mathys, Zhou, Liddelow, and others. Specifically, we gathered the top markers for astrocytes, oligodendrocytes, and microglia from these studies. We then analyzed the relative expression levels of these markers in our own dataset to identify corresponding AD-related cell subtypes mentioned in these publications. This comparative approach allowed us to validate our findings and highlight specific cell types associated with Alzheimer’s disease in our study.

## CRediT authorship contribution statement

**Yunxia Guo:** investigation, methodology, validation, writing-original draft, all authors have read and approved the final manuscript. **Junjie Ma, Ruicheng Qi and Guangzhong Wang:** Data Curation and visualization. **Xiaoying Ma, Jitao Xu, Kaiqiang Ye, Yan Huang and Xi yang:** Writing-review & editing. **Xiangwei Zhao:** Methodology, supervision, resources, project administration and funding acquisition. All authors had read and agreed to the published version of the manuscript.

## Data availability

The single nuclei RNA sequencing data that support the findings of this study have been deposited in the National Center for Biotechnology Information (NCBI) with accession number PRJNA1117576.

## Acknowledgments

This study was supported by the National Natural Science Foundation of China (81827901, 82361138570).

