## Supplementary Material for "snCED-seq: High-fidelity cryogenic enzymatic dissociation of nuclei for single-nucleus RNA-seq of FFPE tissues"

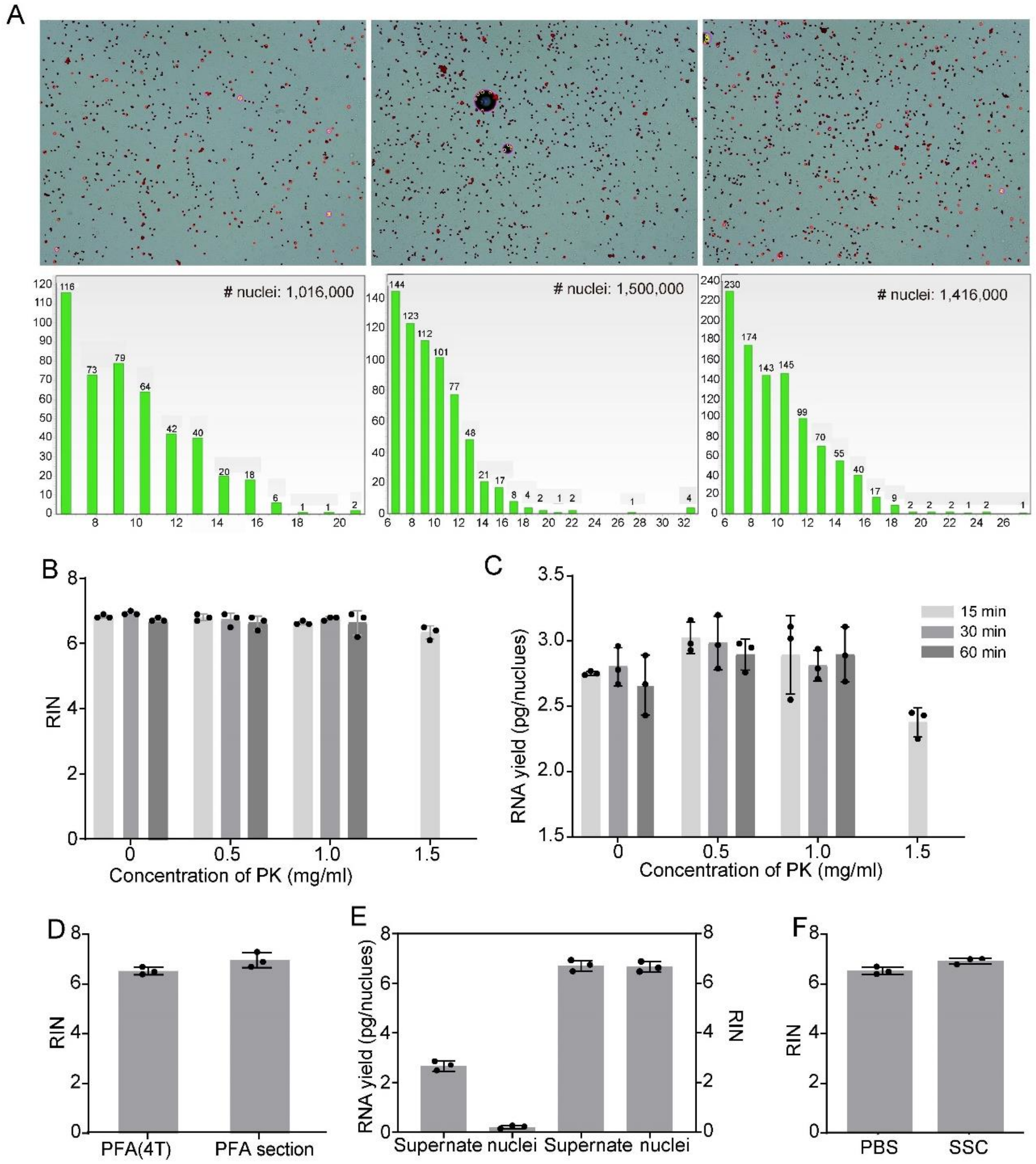


**Fig. S1 The quality of nuclei prepared by CED method was evaluated.** (**A**) The number and diameter distribution of nuclei in triplicate. (**B, C**) Bar plots showing the RIN (B) and yield (C) of total RNA from the bulk nuclei that underwent cross-link heat reversal (15-60 min at 56°C) with proteinase K (0-1.5mg/mL). (**D**) RNA integrity of PFA-fixed brain section and nuclei from a same brain. (**E**) Bar plots showing the yield (Left) and RIN (Right) within the nucleus prepared by HED methods. (**F**) Effect of buffers on RNA integrity of the bulk nuclei. Data are presented as mean ± standard deviation. n=3 technical replicates. Nuclei in all figures were obtained from the CED method, except in figure E.

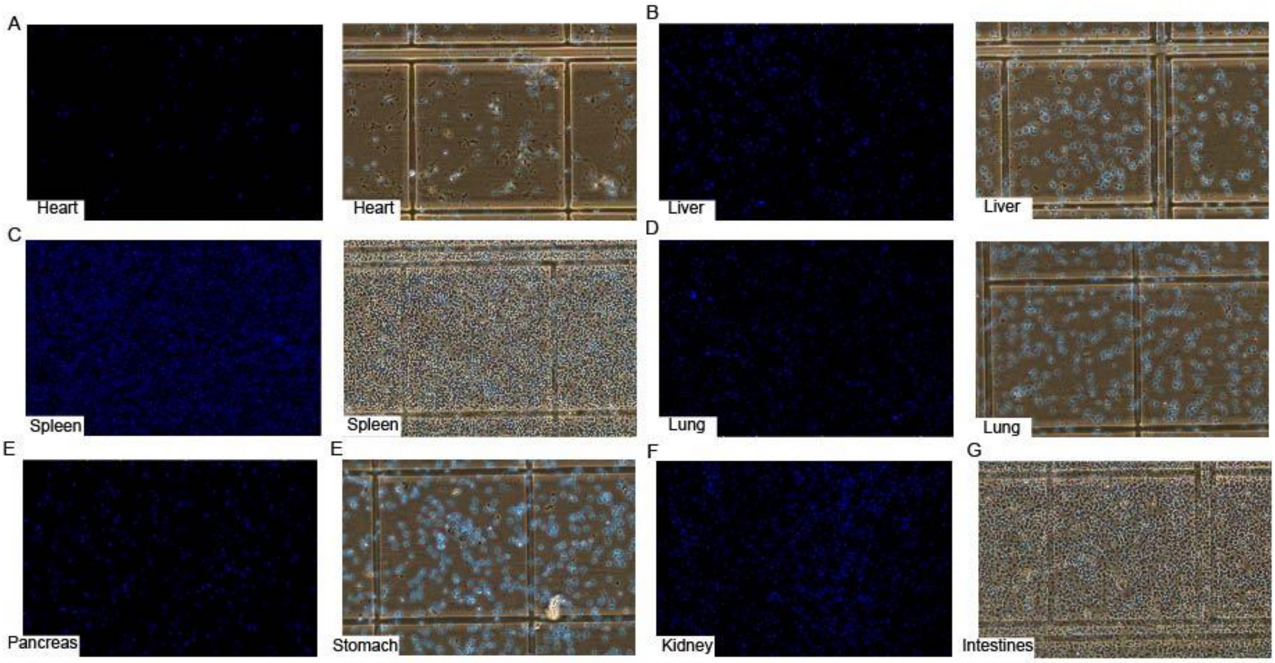


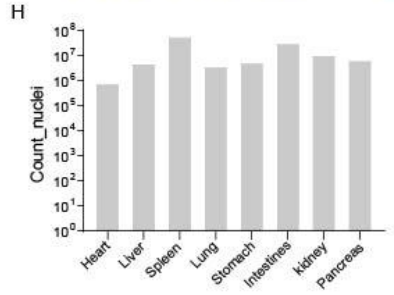


I

| Sample ID | Weight (Before, g) | Weight (After, g) | Weight (sample, g) | NO. of nuclei | Nuclei yield (number/g) |
|---|---|---|---|---|---|
| 1 | 2514.1 | 2543.2 | 29.1 | 3,825,000 | 131,443 |
| 2 | 2536.2 | 2560.7 | 24.5 | 2,925,000 | 119,387 |
| 3 | 2546.3 | 2569.3 | 23 | 2,700,000 | 117,391 |
| 4 | 2532.5 | 2557.7 | 25.2 | 220,000 | 8,730 |
| 5 | 2537.7 | 2564.0 | 26.3 | 187,000 | 7,110 |
| 6 | 2532.5 | 2549.5 | 17 | 250,000 | 14,705 |

**Fig. S2 The morphology of different organs of the nucleus from CED method.** (**A-G**) The microscopic images of nuclei by CED method form heart, liver, spleen, lung, pancreas, stomach, kidney, and small intestine organs, respectively. Scale bar, 50 μm; (**H**) Number of nuclei from A-G; (**I**) Nuclei yield of hippocampus tissue. Nuclei of samples 1-3 and 4-6 were prepared by CED method and commercial nucleus dissociation kit for FFPE sample, respectively.

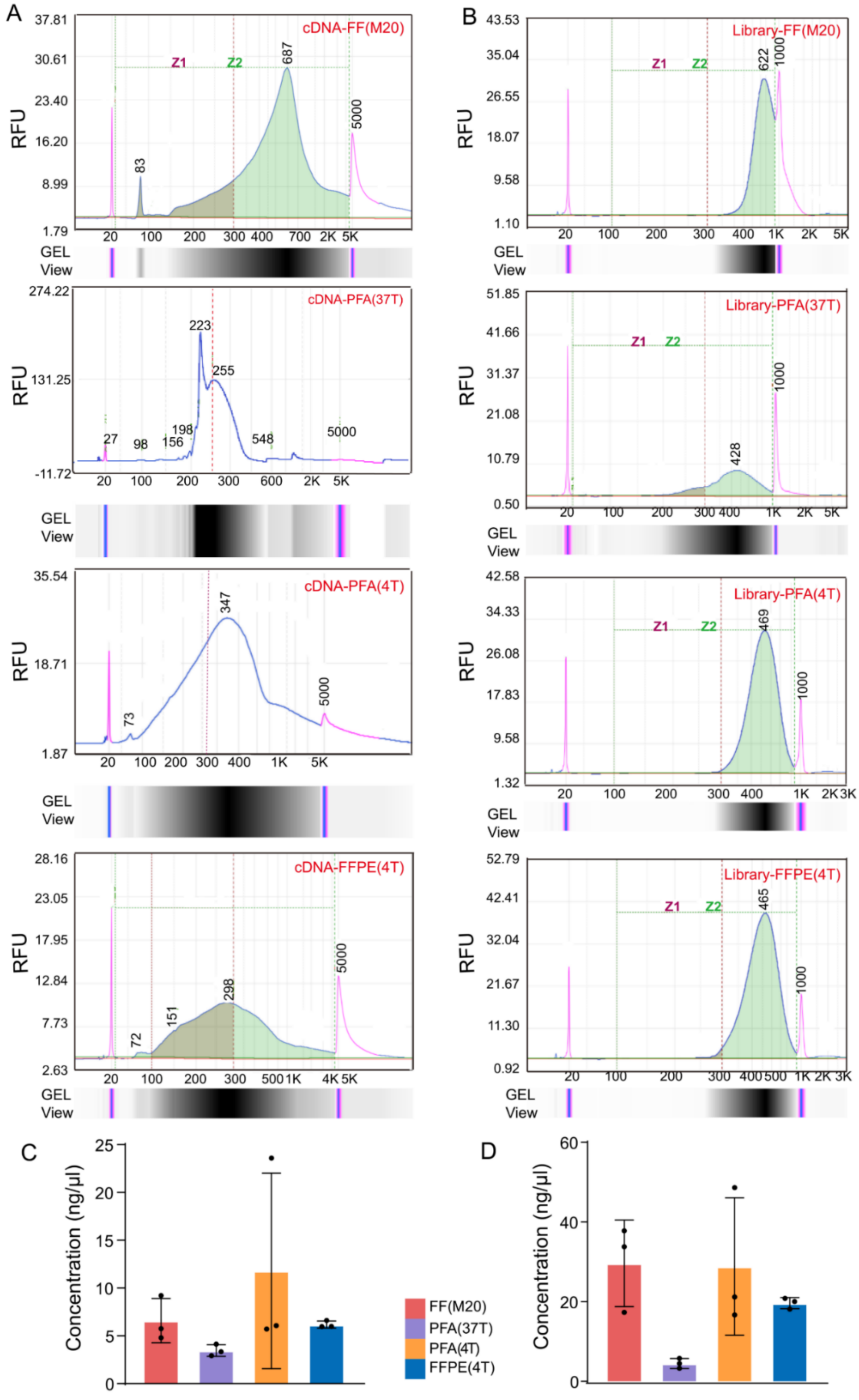


**Fig. S3 The quality of cDNA and libraries generated by snCED-seq from fresh, fixed and FFPE samples.** (**A, B**) Electropherogram of FFPE mouse brain cDNA (A) and library (B) for Qsep100™ DNA Fragment Analyzer. (**C, D**) Bar plots showing the yield of cDNA (C) and library (D) from all group samples.

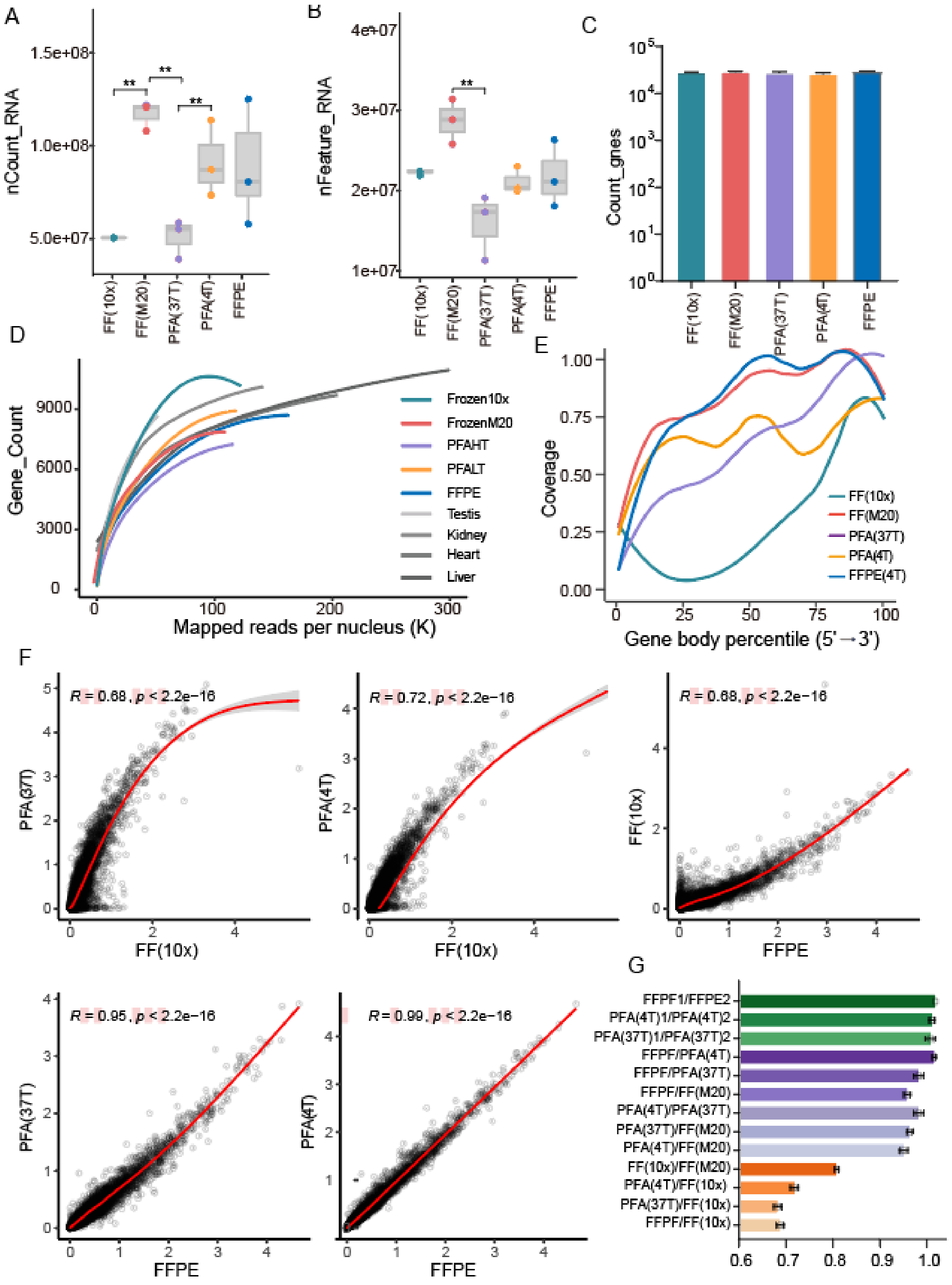


**Fig.S4 Quality analysis of snRNA-seq data.** (**A, B**) Counts of total UMIs (A) and genes (B) from all nuclei of every group samples. (**C**) Counts of total genes detected in different samples. (**D**) Saturation analysis comparison of our brain nuclei and mouse tissues[1] (heart, kidney, testis, and liver) using snRandom-seq. (**E**) Reads distribution along the gene body of all samples. (**F**) The Pearson's correlation coefficient (R) of the normalized gene expressions between samples. Each dot represents the average expression level of a gene. The red line indicates the linear regression line. *p* value (*p*) was computed from two-sided permutation test. (**G**) Bar chart showing the gene expression of correlation between pairwise samples.

A

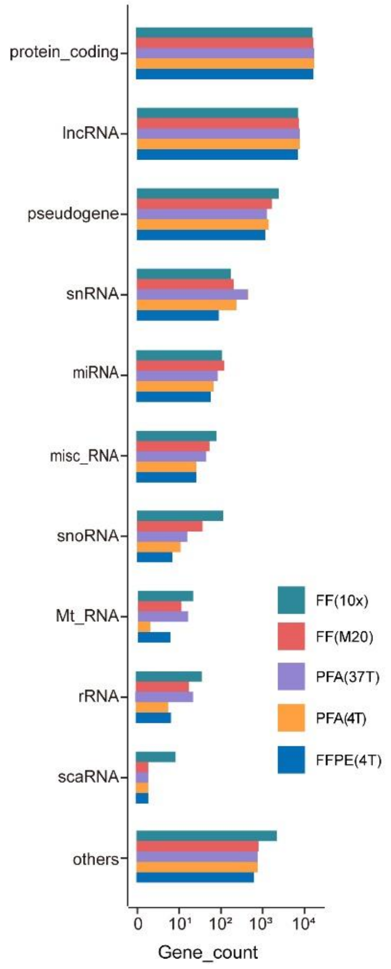


B

| Biotype | snFFPE- seq | snRandom-seq | FF(10x) | FF(M20) | FFPE |
|---|---|---|---|---|---|
| Protein_coding | 12996 | 19564 | 17064 | 17629 | 17736 |
| lncRNA | 921 | 7743 | 7182 | 7420 | 6742 |
| TEC | 9 | 918 | 2255 | 766 | 543 |
| Processed pseudogene | 28 | 2259 | 1967 | 1002 | 470 |
| Unprocessed pseudogene | 3 | 693 | 220 | 407 | 373 |
| transcribed processed pseu. | 4 | 157 | 169 | 135 | 119 |
| snRNA | NA | 580 | 176 | 180 | 88 |
| miRNA | NA | 38 | 109 | 122 | 56 |
| Transcribed unprocessed pseu. | 7 | 180 | 31 | 32 | 36 |
| unitary pseudogene | 1 | 44 | 32 | 34 | 33 |
| misc_RNA | NA | 14 | 80 | 50 | 25 |
| transcribed unitary pseu. | 3 | 24 | 24 | 21 | 22 |
| Pseudogene | 1 | 29 | 21 | 21 | 12 |
| snoRNA | NA | 71 | 116 | 34 | 7 |
| rRNA | NA | 27 | 40 | 18 | 6 |
| IG_C_gene | NA | 10 | 2 | 2 | 5 |
| Mt tRNA | NA | 19 | 19 | 8 | 4 |
| TR_C_gene | NA | 7 | 5 | 3 | 3 |
| TR V gene | NA | 15 | 1 | 1 | 2 |
| IG_V_pseudogene | NA | 12 | NA | NA | 2 |
| IG_V_gene | NA | 10 | NA | NA | 2 |
| TR_V_gene | NA | NA | NA | 2 | 2 |
| scaRNA | NA | 3 | 9 | 2 | 2 |
| Mt_rRNA | NA | 2 | 2 | 2 | 2 |
| translated unprocessed pseu. | 1 | 2 | 1 | 1 | 1 |

**Fig.S5 snRNA-seq data from mouse brains.** (**A**) Biotypes detected from all groups. (**B**) Counts of different biotypes detected in FFPE mouse liver by snRandom-seq, mouse brain by snCED-seq and mouse brain by 10× Genomics.

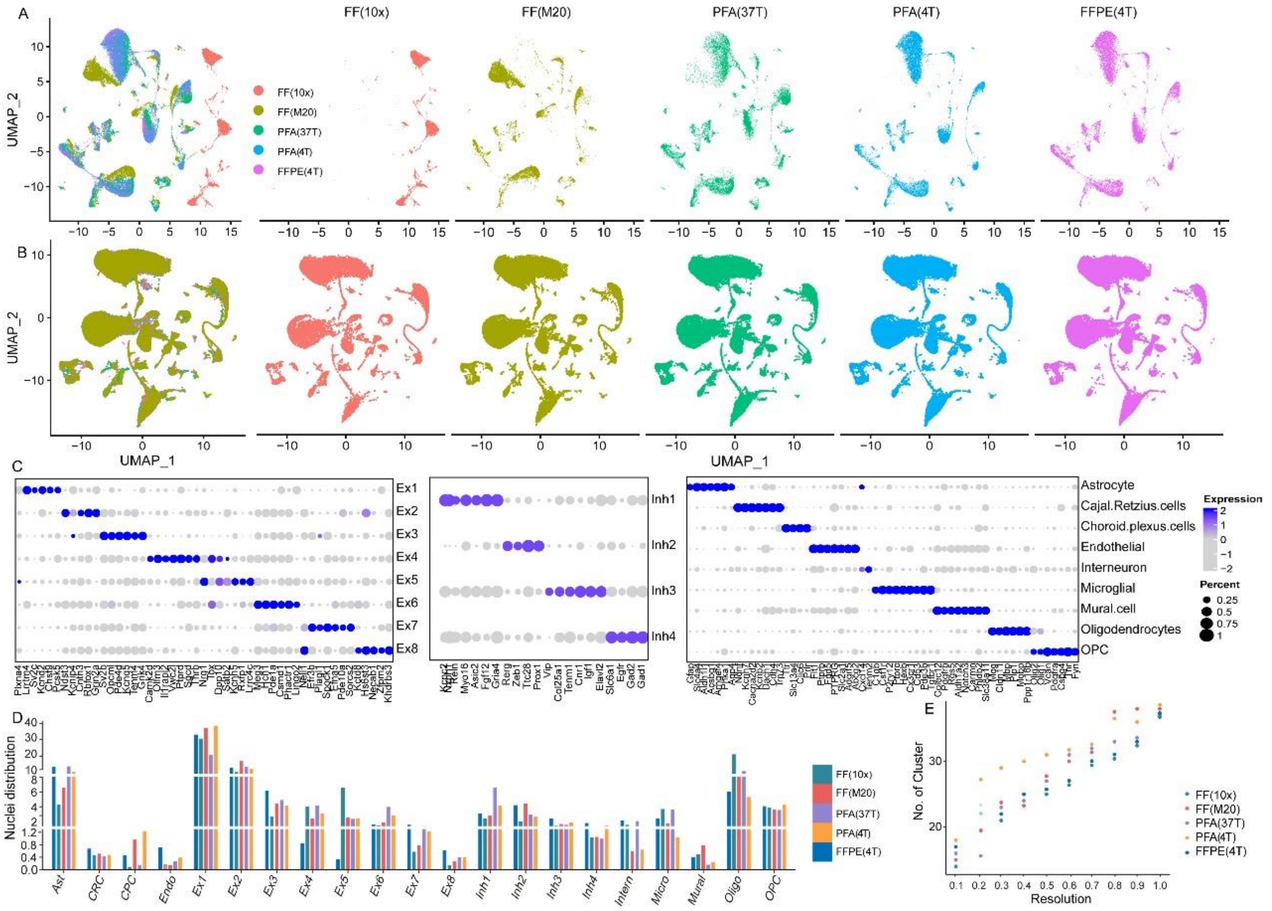


**Fig.S6 Cell type marker genes and assignments. (A, B)** UMAP embedding of single nuclei RNA profiles before (A) and after (B) removing batch effect. The UMAP maps of FF(10×) was significantly different from other samples before removing batch. (**C**) cells and marker genes. Dot plot showing the expression level (color scale) and the percent of cells expressing (dot size) marker genes across all clusters (rows). (**D**) percent of cell types from all groups. (**E**) Number of cell clusters at different resolutions by unsupervised clustering analysis.

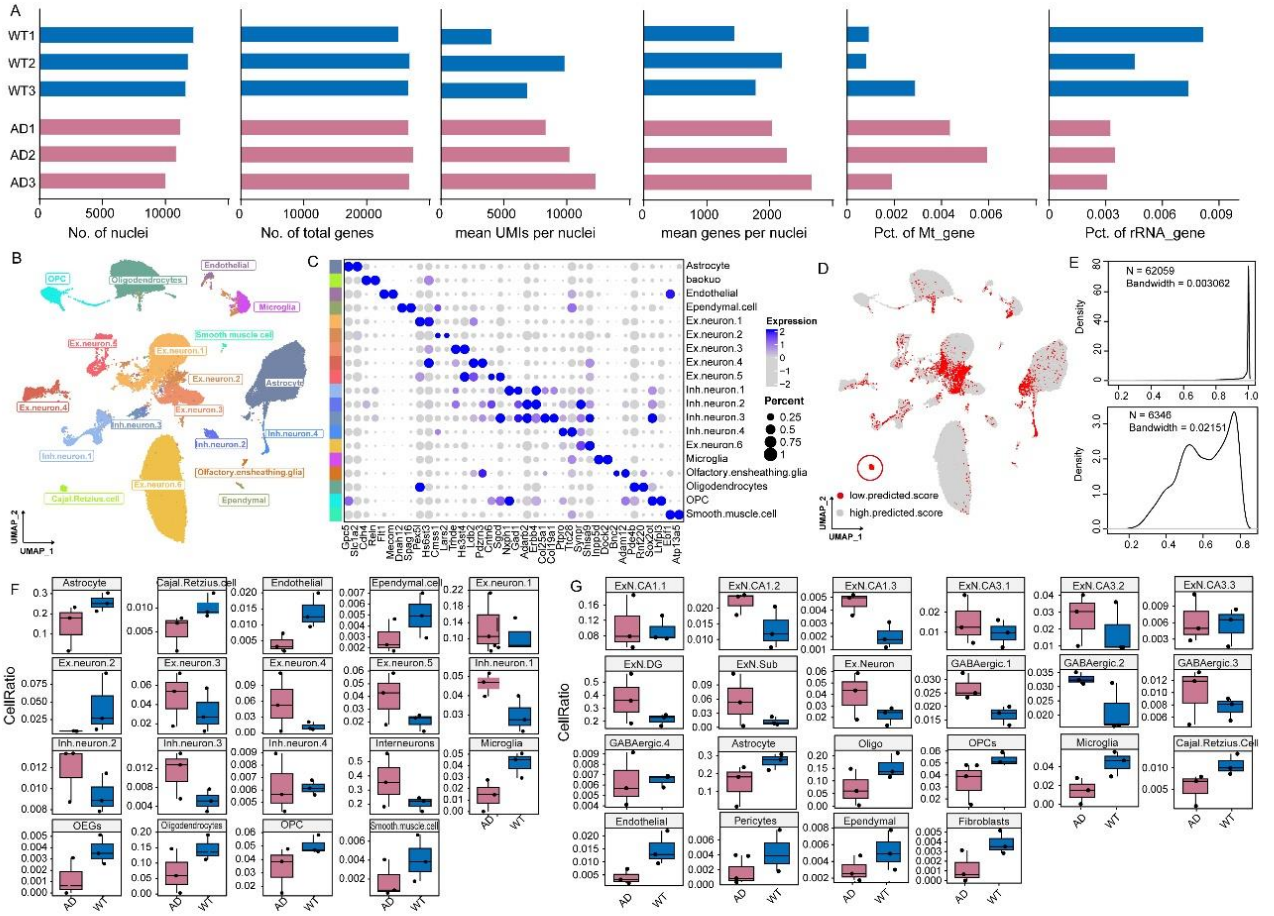

**Fig.S7 Cellular maps of the mouse hippocampus of WT and 5xFAD mice and quality controls.** (**A**) Bar graphs show, from left to right, the distribution of number of nuclei, total genes, transcripts (unique UMIs) per nuclei and genes per nuclei, percent of mitochondrial genes and ribosomal genes detected in WT and AD, respectively. (**B**) UMAP map of AD and WT of snRNA-seq data by unsupervised clustering. (**C**) Cell specificity marker genes in B. (**D**) Cell types predictive analysis between supervised cluster and reference data [2]. Red indicates low predicted and gray indicates high predicted. (E) High prediction (top) and low prediction (bottom) scores for supervised and reference data [2]. (**F, G**) The percent cells of AD and WT in same snRNA-seq data by unsupervised clustering (F) and supervised clustering (G), respectively.

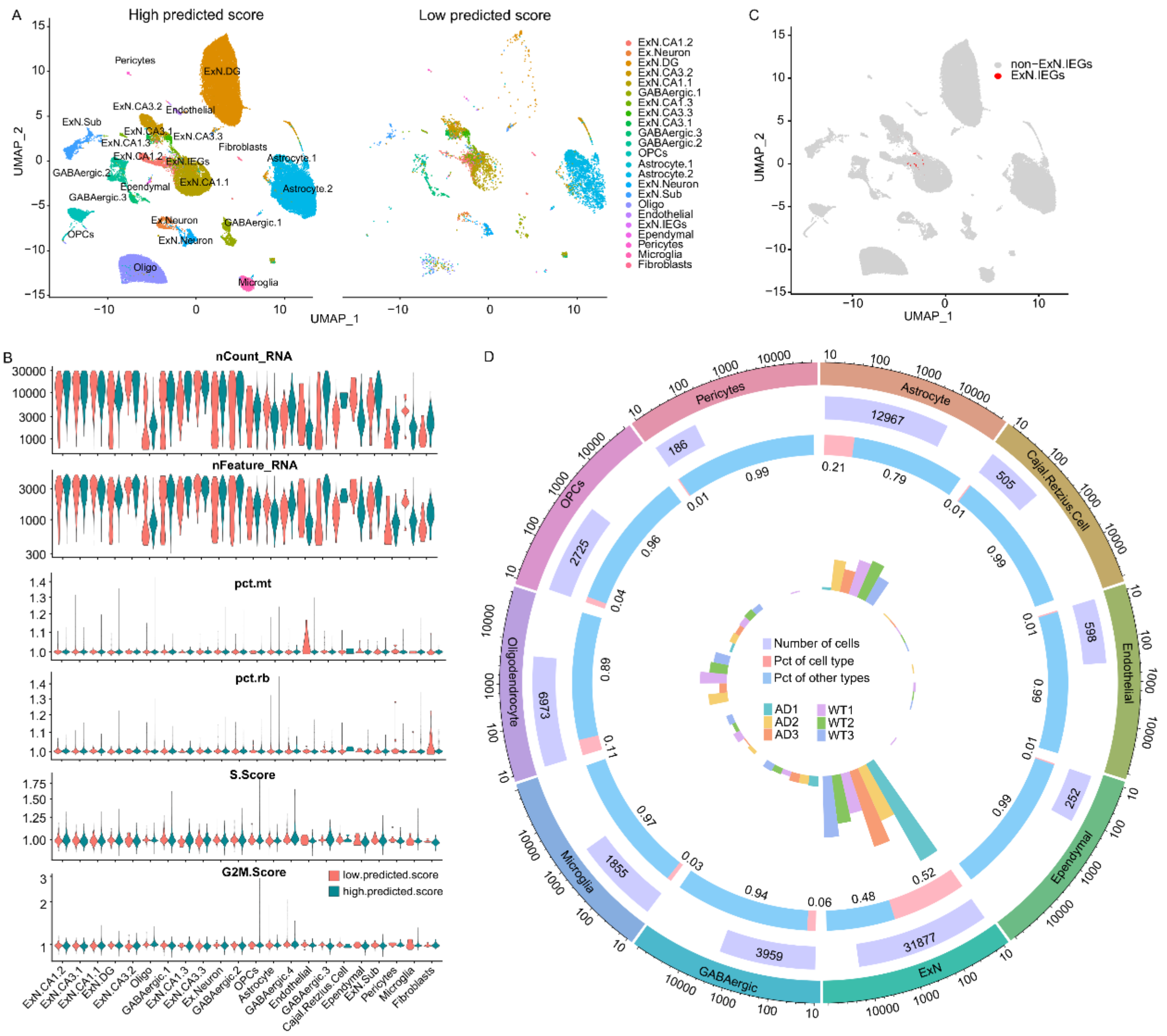


**Fig.S8 Difference analysis of unsupervised clustering and supervised clustering.** (**A**) UMAP atlases of cell types with high (left) and low (right) prediction in supervised clustering. (**B**) Prediction analysis of nCount, nFeature, percent of mitochondrial and ribosomal genes, and cell cycle of cell types. (**C**) The distribution of ExN.IEG cells in UMAP atlas. (**D**) Infographic of cell types in AD and WT. Cell types, the number of nuclei, the percentage of cells, expressed in different colors.

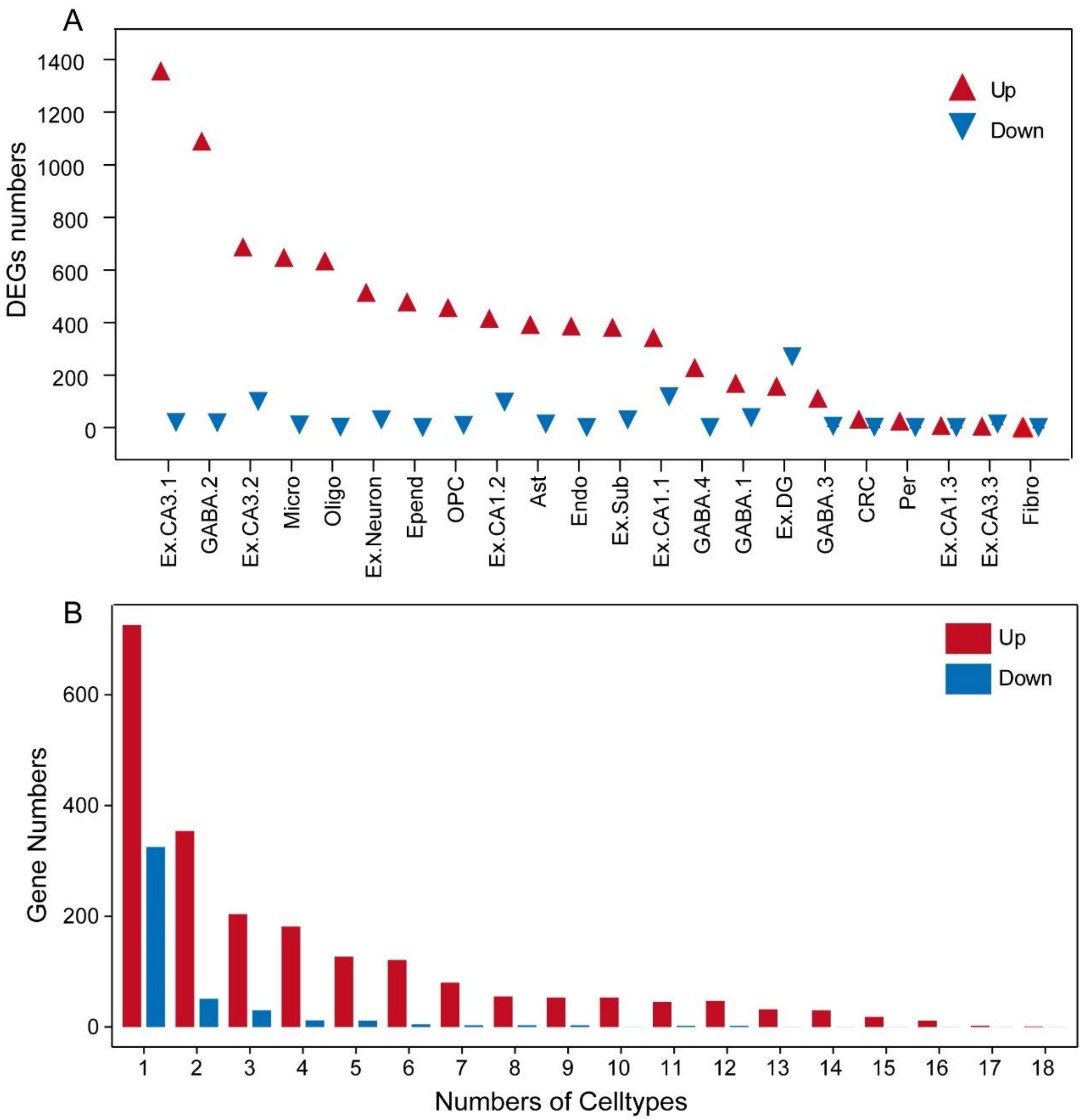


**Fig.S9 Cell-type-specific gene-expression changes in AD.** (**A**) Progressive changes in DEG counts for each cell type in AD and WT. (**B**) The number of up- or down-regulated genes per cell type was detected. Red and blue respectively up and down DEGs.

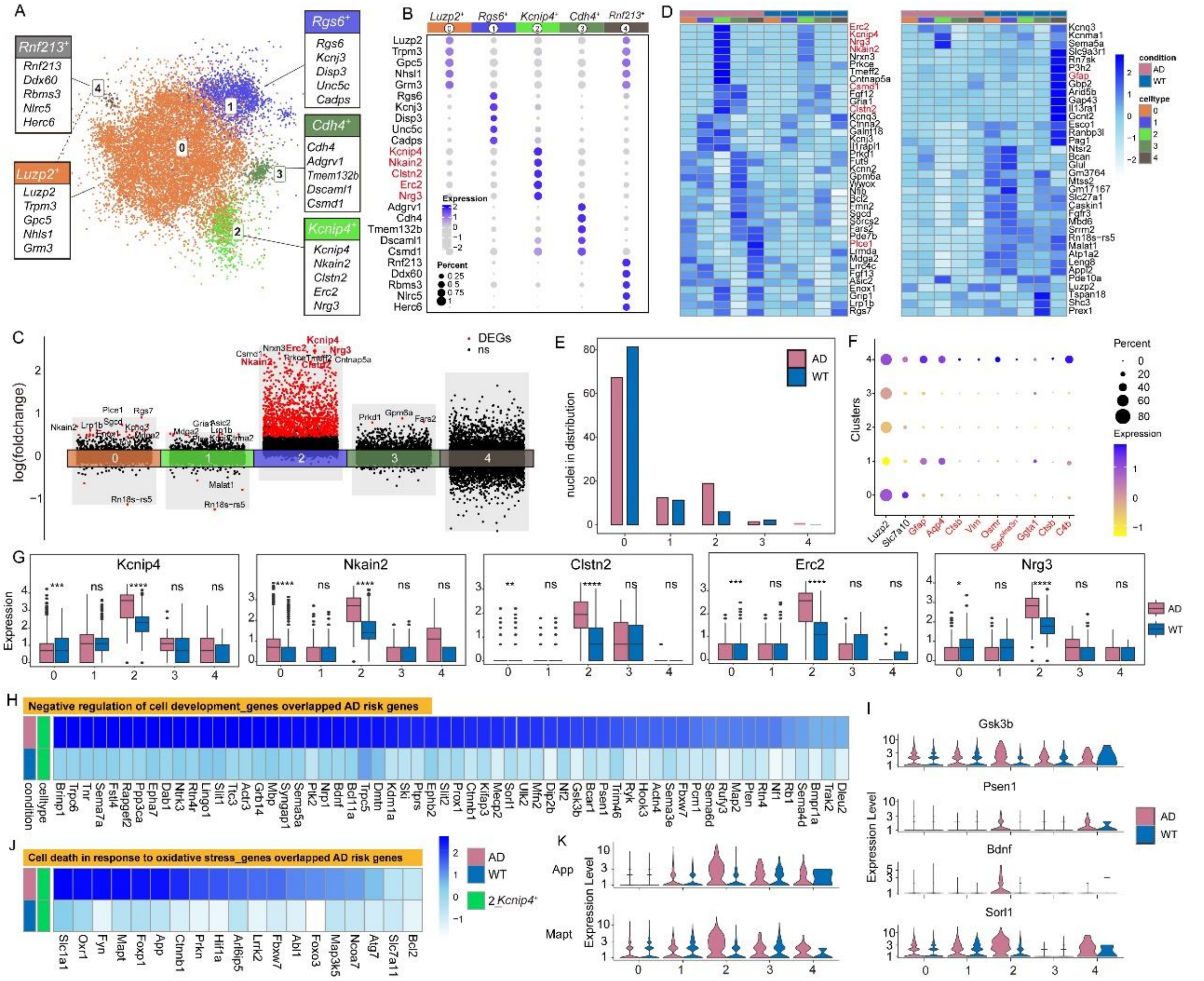

**Fig.S10 Characterization of the DAAs in AD.** (**A**) UMAP plot of re-clustered astrocytes identifying 5 sub-clusters. n = 3 biologically independent mouse brain samples per genotype; 2,840 total astrocytes nuclei. (**B**) The average gene expression of top DEGs in the astrocyte sub-clusters. Expression level (color scale) of marker genes across clusters and the percentage of cells expressing them (dot size). (**C**) Volcano plots showing significant DEGs (fold change > 1.5, adjusted $p < 0.05$, Bonferroni correction) in astrocytes of 5xFAD versus WT. (**D**) Average scaled expression of the top-10 (Left) upregulated and (Right) downregulated disease-specific DEGs split by cluster. Color scheme shows row max and row min, which represents relative expression of each gene among all sub-clusters. (**E**) Bar graph showing the frequency of each astrocytes sub-cluster in AD and WT. (**F**) DAA genes specificity high expression in AST4. (**G**) Top upregulated DEGs (DAA genes) of AD were specifically highly expressed in AST2. (**H-K**) The genes associated with disease-related function or pathway were highly expressed in AST2.

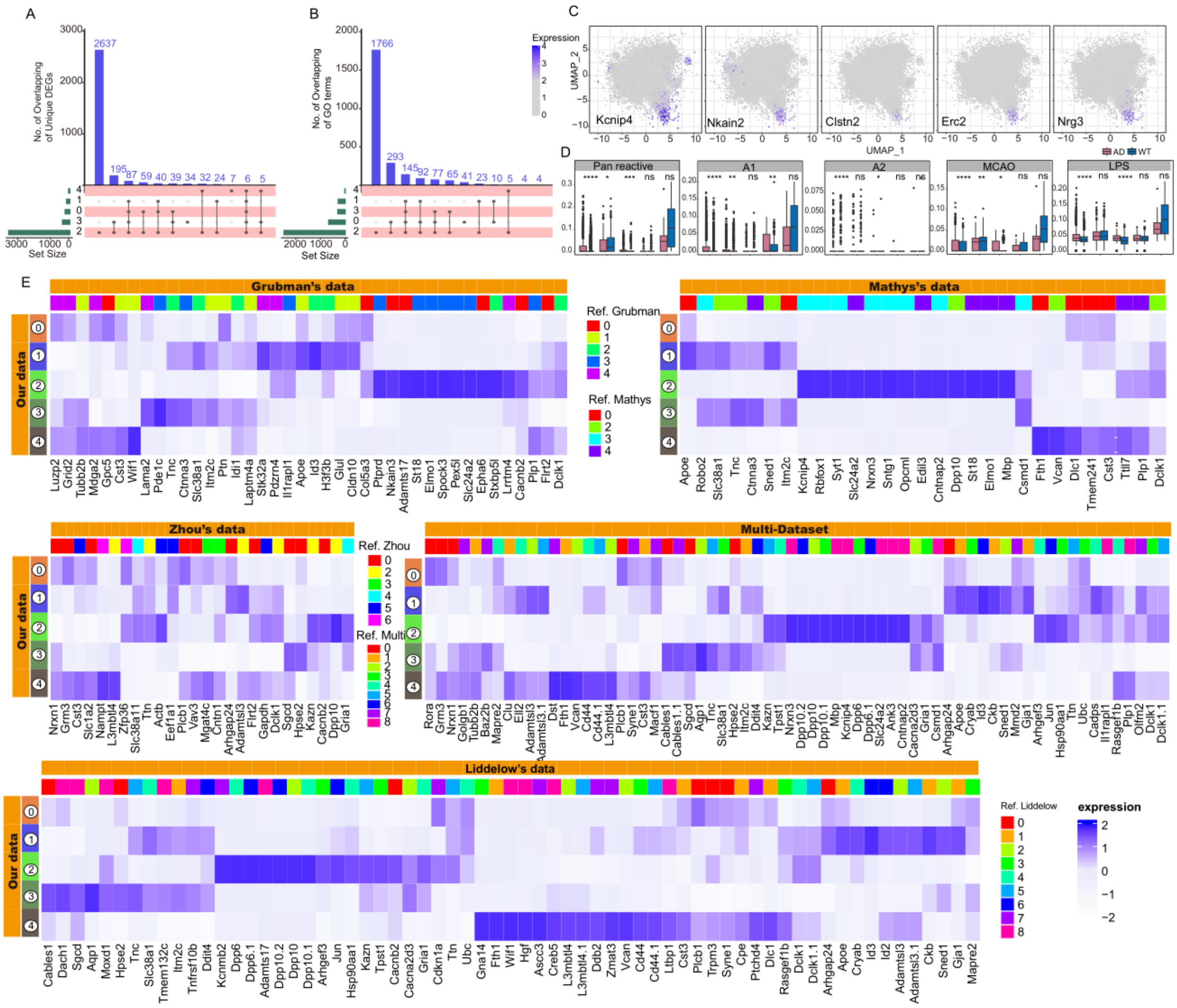

**Fig.S11 Integration of astrocytes from multiple datasets reveals consistent identification of astrocytes subtypes.** (**A, B**) UpSetR plots highlighting upregulated DEGs (A) and upregulated GO terms (B) that are unique to or shared between clusters. Bars show number of DEGs per cluster (colored at left). Lines between clusters highlight shared DEGs. (**C**) Expression of marker genes across AST2. UMAP embedding of single nuclei RNA profiles from hippocampi of WT and 5xFAD mice, colored by expression levels of marker genes. (**D**) Signatures of reactive astrocytes found in AD. Box-plot showing the distribution of expression scores for previously defined signature genes for (left) pan-reactive, A1/A2-reactive, ischemia-associated (middle cerebral artery occlusion, MCAO) reactive and inflammation-associated (lipopolysaccharides, LPS) astrocytes across clusters. $p > 0.05$, ns. (**E**) Average scaled expression heatmap of the top 5 cluster-enriched/unique transcripts per cluster in the published snRNA-seq dataset (Grubman [3], Mathys [4], Zhou [5], Liddelow [6], and integrated data) for astrocytes in AD in our data.

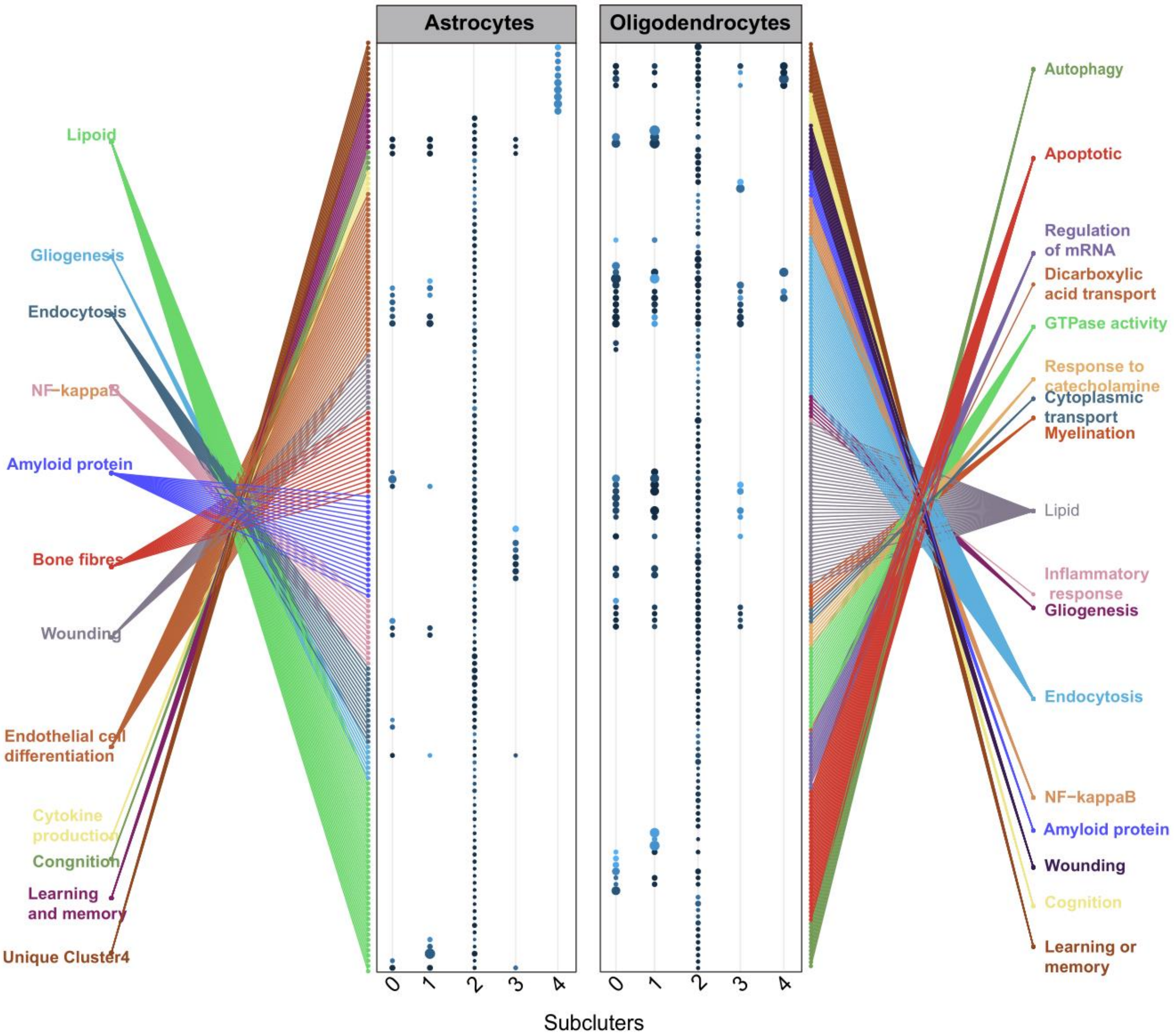


**Fig.S12 Astrocytes (A) and oligodendrocytes (B) subclusters enrichment the reported function and pathways associated with AD.**

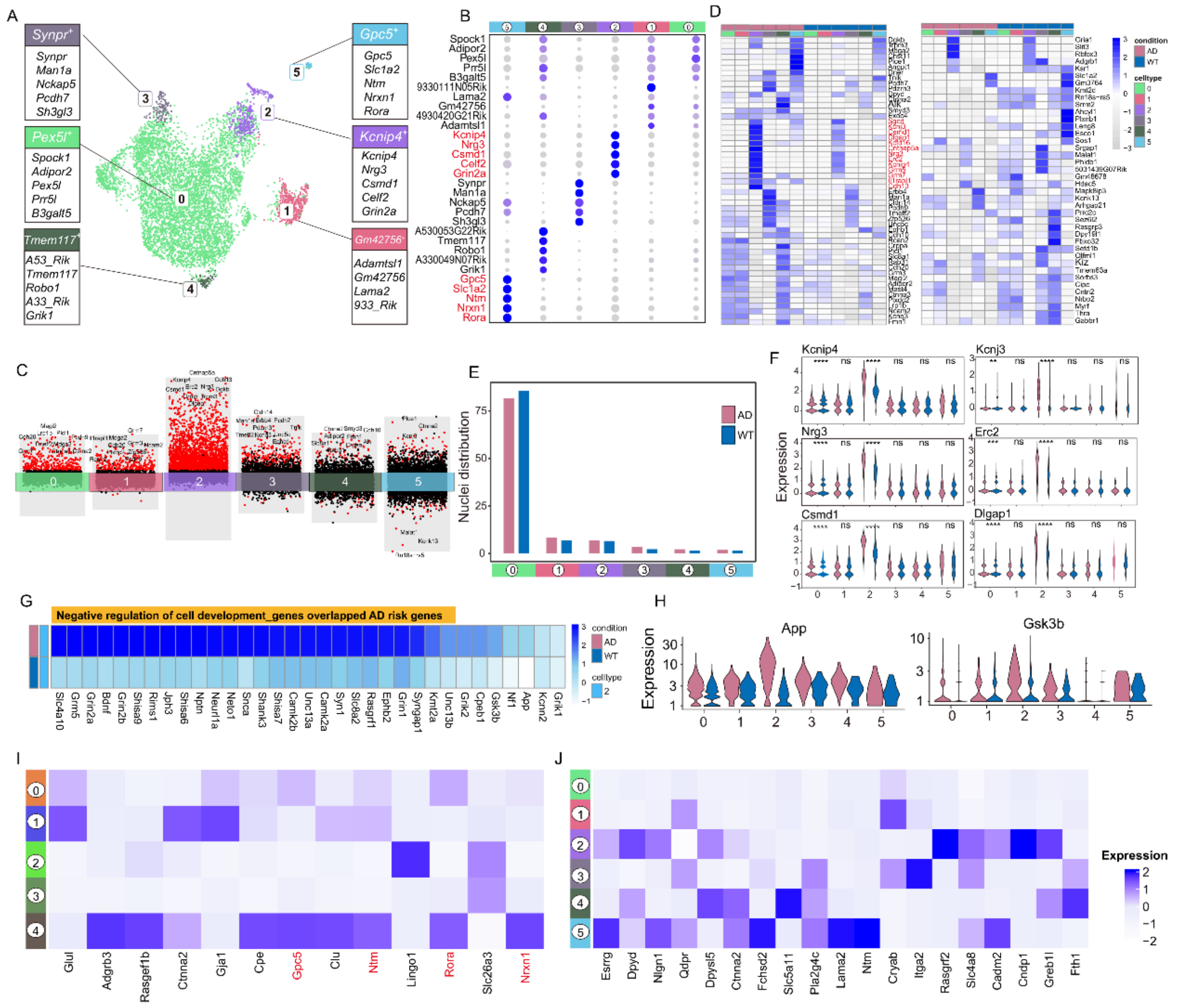


**Fig.S13 Oligodendrocytes have cluster-specific transcriptomic changes in Alzheimer's disease.** (**A**) UMAP plot of re-clustered oligodendrocytes identifying 6 sub-clusters. (**B**) Marker genes of oligodendrocytes states. Expression level (color scale) of marker genes across clusters and the percentage of cells expressing them (dot size). (**C**) Volcano plots showing significant DEGs in oligodendrocytes subclusters of AD versus WT. (**D**) Average scaled expression of the top-10 (Left) upregulated and (Right) downregulated disease-specific DEGs split by cluster. (**E**) Bar graph showing the frequency of each oligodendrocytes sub-cluster in AD and WT. (**F**) Top upregulated DEGs of AD in E were specifically highly expressed in Oligo2. (**G, H**) The genes associated with disease-related function were highly expressed in Oligo2 and Oligo5 of AD. (**I, J**) Heat map of amyloid-related cell type-specific genes expression in the astrocyte (I) and oligodendrocyte subcluster (J).

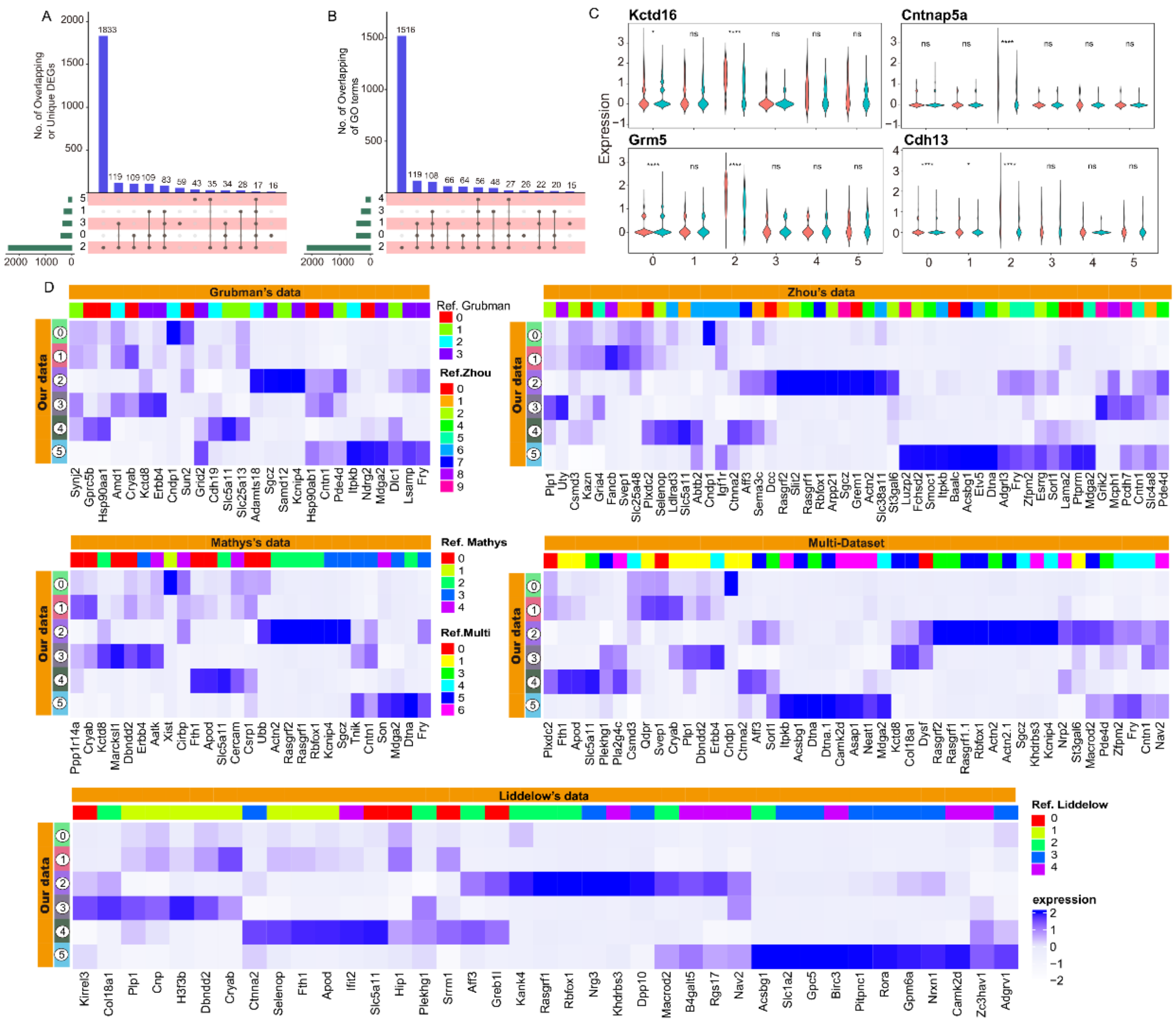


**Fig.S14 Integration the multiple datasets reveal consistent identification of oligodendrocytes subtypes.** (**A, B**) UpSetR plots highlighting upregulated DEGs (A) and upregulated GO terms (B) that are unique to or shared between clusters. (**C**) Expression of marker genes across Oligo2. (**D**) Average scaled expression heatmap of the top 5 cluster-enriched/unique transcripts per cluster in the published snRNA-seq dataset (Grubman [3], Mathys [4], Zhou [5], Liddelow [6], and integrated data) for oligodendrocytes in AD in our data.

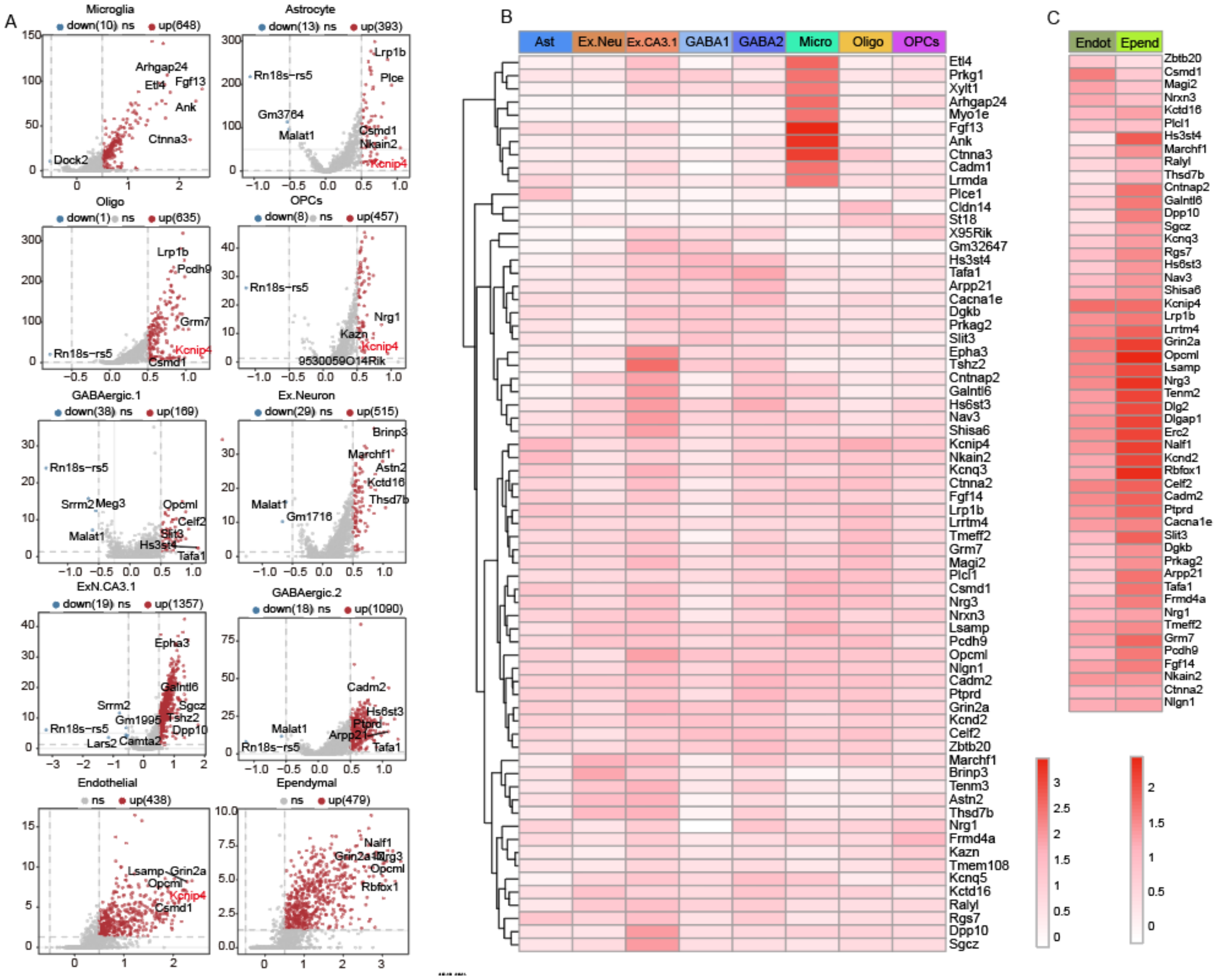

**Fig.S15 Disease-related cell types share transcriptional similarities.** (**A**) Volcano plots showing significant DEGs in AD-related cells of AD versus WT. (**B, C**) Heat maps of top10 DEGs of cell types in A.
